# Relationship between input connectivity, morphology and orientation tuning of layer 2/3 pyramidal cells in mouse visual cortex

**DOI:** 10.1101/2020.06.03.127191

**Authors:** Simon Weiler, Drago Guggiana Nilo, Tobias Bonhoeffer, Mark Hübener, Tobias Rose, Volker Scheuss

## Abstract

Neocortical pyramidal cells (PCs) display functional specializations defined by their excitatory and inhibitory circuit connectivity. For layer 2/3 (L2/3) PCs, little is known about the detailed relationship between their neuronal response properties, dendritic structure and their underlying circuit connectivity at the level of single cells. Here, we ask whether L2/3 PCs in mouse primary visual cortex (V1) differ in their functional intra- and interlaminar connectivity patterns, and how this relates to differences in visual response properties. Using a combined approach, we first characterized the orientation and direction tuning of individual L2/3 PCs with *in vivo* 2-photon calcium imaging. Subsequently, we performed excitatory and inhibitory synaptic input mapping of the same L2/3 PCs in brain slices using laser scanning photostimulation (LSPS).

Our data from this structure-connectivity-function analysis show that the sources of excitatory and inhibitory synaptic input are different in their laminar origin and horizontal location with respect to cell position: On average, L2/3 PCs receive more inhibition than excitation from within L2/3, whereas excitation dominates input from L4 and L5. Horizontally, inhibitory input originates from locations closer to the horizontal position of the soma, while excitatory input arises from more distant locations in L4 and L5. In L2/3, the excitatory and inhibitory inputs spatially overlap on average. Importantly, at the level of individual neurons, PCs receive inputs from presynaptic cells located spatially offset, vertically and horizontally, relative to the soma. These input offsets show a systematic correlation with the preferred orientation of the postsynaptic L2/3 PC *in vivo*. Unexpectedly, this correlation is higher for inhibitory input offsets within L2/3 than for excitatory input offsets. When relating the dendritic complexity of L2/3 PCs to their orientation tuning, we find that sharply tuned cells have a less complex apical tree compared to broadly tuned cells. These results indicate that the spatial input offsets of the functional input connectivity are linked to orientation preference, while the orientation selectivity of L2/3 PCs is more related to the dendritic complexity.

## Introduction

A fundamental question in neuroscience is how neural activity during sensory processing or behavior arises from underlying principles at the circuit, cellular and synaptic level. One aspect of this is to understand the relationship between activity patterns and synaptic connectivity within the neuronal circuit. The neocortex of mammals by and large displays a universal organization at the circuit level, with only limited variations between cortical areas and species (reviewed in Douglas & Martin, 2004; Harris & Shepherd, 2015). This so-called canonical circuit has been extended by additional intra- and interlaminar connections, the wiring of interneurons and specific subclasses of principal cells, as well as the identification of subnetworks of preferentially connected sets of neurons across laminae (Harris & Mrsic-Flogel, 2013). Advances in experimental techniques over the last years have confirmed previously predicted principles of neuronal organization: 1) Neurons inherit response properties from their input neurons to some extent, and reciprocally connected cells amplify these cortical responses (Wertz et al. 2015; Ko et al. 2011; Lien and Scanziani 2013). 2) Cell types with different morphologies, electrophysiological properties and connectivities display different response characteristics (Vélez-Fort et al. 2014; Kim et al. 2015). 3) Inhibition contributes to and sharpens the tuning properties of neurons (Wilson, Scholl, and Fitzpatrick 2018; Liu et al. 2011). However, whether and how specific connectivity motifs at the circuit level relate to particular stimulus response properties remains unclear. Here, we address circuit mechanisms of cortical function by exploring the relationship between the circuit connectivity motifs of individual neurons and their specific response properties in mouse visual cortex. The retinotopic organization of the visual cortex permits relating the spatial arrangement of input neurons to the spatio-temporal dynamics in visual space of preferred and non-preferred visual stimuli, thereby allowing for the inference of functional circuit mechanisms. Basic principles of circuit models suggested for visual feature selectivity are spatial sampling biases and spatiotemporal offsets in the integration of stimuli across visual space. For example, orientation tuning is thought to arise from selectively combining inputs that respond to stimuli at spatial locations laying along the orientation of the preferred stimulus (Hubel & Wiesel, 1962; Chapman, Zahs, & Stryker, 1991; Alonso & Reid, 1995).

We focused on L2/3 pyramidal cells because they are at the core of cortical processing, in between the input and output layers. Although L2/3 is subject to many *in vivo* studies on neuronal function and plasticity, there is still little mechanistic insight into how exactly response properties arise from intracortical connectivity. For pyramidal cells in L5 and L6, a correspondence between morphological, electrophysiological and functional response characteristics has been established and led to the distinction of several cell types (Vélez-Fort et al. 2014; Kim et al. 2015). Whether pyramidal cells in L2/3 also consist of different subpopulations has not yet been reported. With respect to circuit mechanisms, it was shown for example in ferret and mouse visual cortex, at the postsynaptic dendritic level, that the sum of excitatory inputs predicts the preferred orientation of L2/3 pyramidal cells (Chen et al. 2013; Wilson et al. 2016), and that the selectivity of orientation tuning depends on functional clustering of synaptic inputs (Wilson et al. 2016). However, the relationship between neuronal morphology, circuit connectivity and functional response properties of L2/3 cells has not been systematically analyzed so far.

To address this question, we used a combined structure-connectivity-function analysis. We first characterized the tuning properties of individual L2/3 PCs in V1 using *in vivo* two photon calcium imaging. We then reidentified the same L2/3 PCs in acute brain slices and mapped their intra- and interlaminar excitatory and inhibitory inputs using laser scanning photostimulation (LSPS) by UV glutamate uncaging (Weiler et al. 2018). Simultaneously, we filled the cells with Alexa-594 and reconstructed their dendritic tree. We found that the intra- and interlaminar inputs are diverse among L2/3 PCs, with mostly spatially balanced excitation and inhibition at the population level. On a single cell level, we found vertical and horizontal offsets of excitatory and inhibitory inputs. These input offsets were directly related to the preferred orientation of the postsynaptic cell, and L2/3 PCs with the largest difference between these offsets had preferred orientations that were orthogonal to each other. While the preferred orientation was related to the cell’s synaptic input, the tuning selectivity was directly related to the apical but not the basal tree complexity. L2/3 PCs with a less complex apical tree had a higher orientation selectivity compared to cells with a more complex apical tree.

## Results

### Visual response properties and neuronal circuit connectivity of the same neurons

For studying the relationship between neuronal circuit connectivity motifs and sensory processing, we recorded both visual response properties and synaptic inputs in the same pyramidal cells (PCs) in mouse visual cortex (Fig. 1A, Weiler et al., 2018). We sparsely co-expressed GCaMP6m and mRuby2 by viral transduction in the binocular zone of primary visual cortex (bV1) and performed functional and structural imaging. We recorded neural activity in individual PCs of L2/3, both in the dark and evoked by eye-specific visual stimuli (Fig. 1C). One day after characterization of visual response properties, we prepared acute coronal slices containing the imaged PCs and mapped local functional inputs by LSPS (Fig. 1D). Individual PCs that had been imaged *in vivo* were identified by comparing relative positions, morphological details, and anatomical landmarks such as blood vessels. This was done in both, the top view of an image stack acquired in the slice by structural 2-photon imaging, and the corresponding side view in an image stack acquired during *in vivo* imaging (Fig. 1C, D). We recorded EPSCs and IPSCs from identified PCs evoked by LSPS via UV-glutamate uncaging for mapping their excitatory and inhibitory presynaptic neurons in the different cortical layers of the slice (16×16 stimulus grid, 69 µm spacing, Supplementary Fig. 2-3, Callaway and Katz 1993; Shepherd, Pologruto, and Svoboda 2003). In addition, the neuron was filled with Alexa-594, and its morphology was assessed by 2-photon imaging, allowing structural analysis of its dendritic tree. In total, we recorded the local excitatory and inhibitory synaptic input of 147 L2/3 PCs. For 70 of these cells we also characterized the visual tuning properties *in vivo* (pial depths between 150 - 350 µm; Fig. 1B). In addition, we obtained the input maps together with the morphology for 97 of all cells (n=147), and the visual response properties, input maps and morphology from 32 of all cells (Fig. 1B, overview of all cells in Supplementary Fig.1).

**Figure 1:**
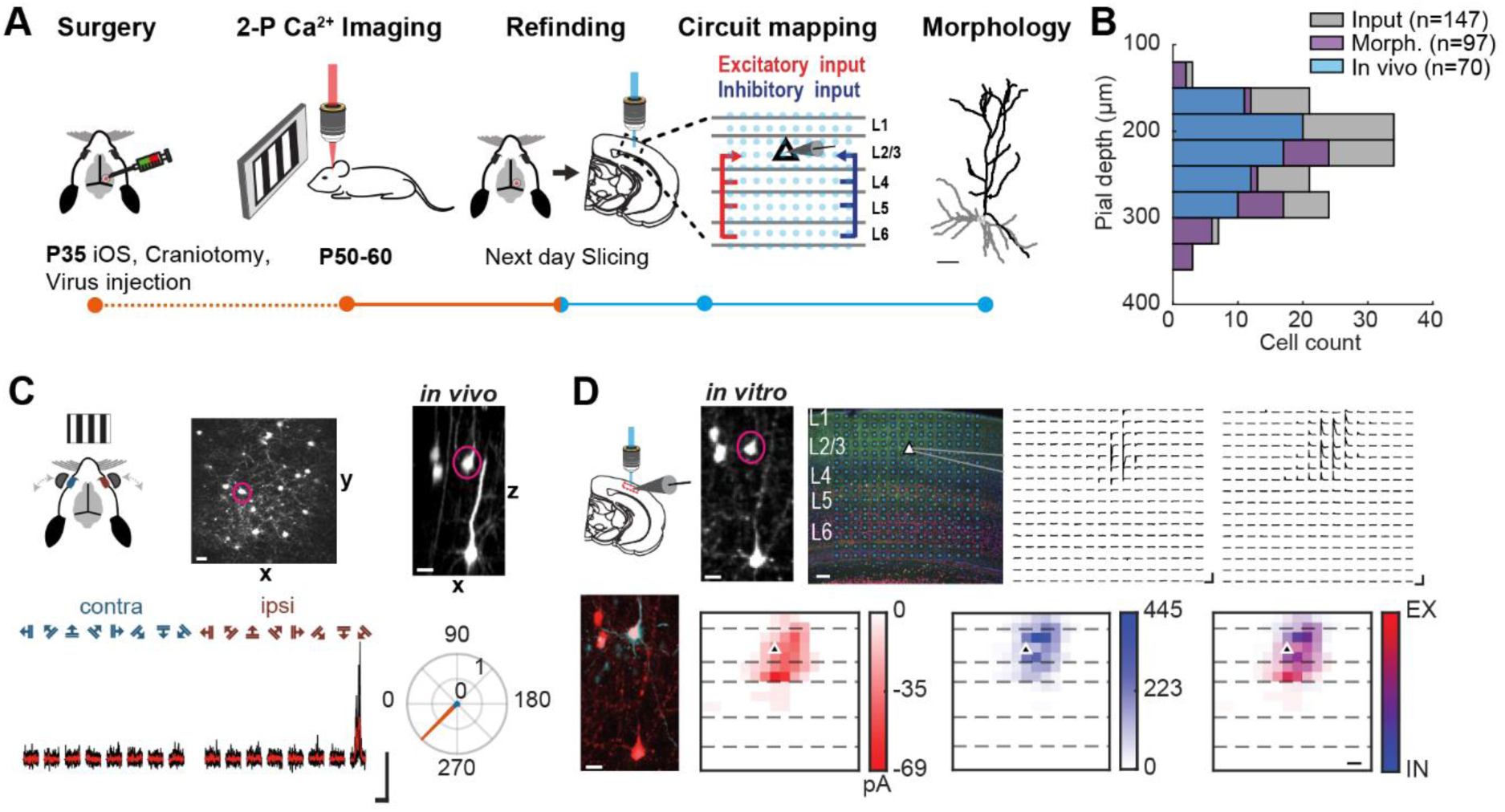
*In vivo* / *in vitro* approach to measure visual response properties and corresponding laminar excitatory and inhibitory inputs of L2/3 pyramidal cells in V1. **A** Experimental *in vivo* / *in vitro* pipeline: GCaMP6m and mRuby2 were expressed in bV1 and the eye-specific orientation/direction tuning of L2/3 PCs was characterized using *in vivo* 2-photon calcium imaging. Subsequently, L2/3 PCs were re-found in acute brain slices and the laminar excitatory and inhibitory inputs were mapped using LSPS by UV glutamate uncaging. Additionally, L2/3 PCs were filled with Alexa-594 to reconstruct their dendritic morphology. Reconstructed L2/3 PC dendritic morphology is from cell shown in C, D (scale bar: 50 µm). **B** Distribution of distances to the pial surface of recorded neurons within L2/3. In total, the laminar input of 147 L2/3 PCs was characterized (grey). Subsets of L2/3 PCs that were in addition morphologically reconstructed and/or functionally characterized are indicated with different colors (magenta and blue, respectively). **C, D** Example of an *in vivo* / *in vitro* characterized cell. **C** Independent eye stimulation paradigm (top, depicted in schematic). The L2/3 PC of interest is marked by a circle in the top and side view of the structural image stack obtained *in vivo* (top, maximum intensity projections; scale bar: 25 µm). Calcium transients of the cell in response to ipsi- or contralateral eye stimulation with drifting gratings of 8 orientations (bottom). Individual calcium transients are shown in black; average in red (scale bars: ΔR/R_0_=200%, 10 s). Polar plot of peak-normalized directional responses to contra- and ipsilateral eye stimulation (blue and red, respectively). **D** Circuit mapping *in vitro*. The L2/3 PC depicted in C is marked by a circle in the *in vitro* side view (top, maximum intensity projection; scale bar: 25 µm). Stimulation grid (blue dots) on brain slice with schematic patch pipette on L2/3 PC. Histological labels mark neocortical layers: L2/3, Calbindin (green); L5/6, CTIP2 (red). All cells stained with DAPI (blue, scale bar: 100 µm). Excitatory currents (cell clamped to −70 mV) and inhibitory currents (cell clamped to 0 mV; scale bars: 250 pA, 500 ms) evoked at corresponding stimulus grid locations. The cell was filled with Alexa-594 (bottom, cyan). Reconstructed dendritic morphology is shown in A. Pixel-based excitatory (red) and inhibitory (blue) input maps represented with color-coded response amplitudes and overlap of both (scale bar: 100 µm).

### Organization of local excitatory and inhibitory inputs to L2/3 pyramidal cells

To better understand the basic cortical wiring diagram of L2/3 PCs in bV1, and to assess the spatial relationship between their excitatory and inhibitory inputs, we first explored the organization of the input maps. Since we recorded excitatory and inhibitory input in the same cells, we were able to assess this relationship on a cell to cell basis. We found that input maps showed diverse laminar and horizontal synaptic input distributions (Supplementary Fig.1). Figure 2A shows examples of excitatory (red) and inhibitory (blue) input maps for three PCs. For further quantification, we aligned the peak-normalized inputs to the medial-lateral axis (Fig. 2A) and computed the input fraction per stimulus row and column. We excluded any excitatory input from L1 since neurons with somata in L2/3-L5 and apical tuft dendrites in L1 also fired action potentials when their tufts were stimulated in L1 (see Methods, Supplementary Fig. 3). Vertically, most excitatory and inhibitory input arose from L2/3, less from L4 and little from L5 (Fig. 2B, left). On average, L2/3 PCs received slightly more inhibition than excitation from L2/3 itself (Fig. 2B, right). At the level of individual cells, the layer-by-layer excitatory and inhibitory input was balanced only for a minority of cells. Despite the wide distribution of excitation and inhibition at the single cell level, a significant number of cells received stronger inhibition than excitation from L2/3 (Fig. 2D, top, Wilcoxon signed-rank, p<0.001). In contrast, the vast majority of cells received stronger excitation than inhibition from both L4 and L5 (Fig. 2D, middle and bottom, Wilcoxon signed-rank, p<0.001). Horizontally, excitatory and inhibitory input was on average centered on the soma location (Fig. 2C). Interestingly, inhibitory input was more concentrated proximal to the soma and excitatory input dominated the more distal regions (Fig. 2C, right panels). This observation was most prominent in L4 and L5, where for the majority of cells, the sources of excitatory input extended further than the sources of inhibition when comparing their horizontal extent (Fig. 2E, middle and bottom, Wilcoxon signed-rank, p<0.001).

**Figure 2:**
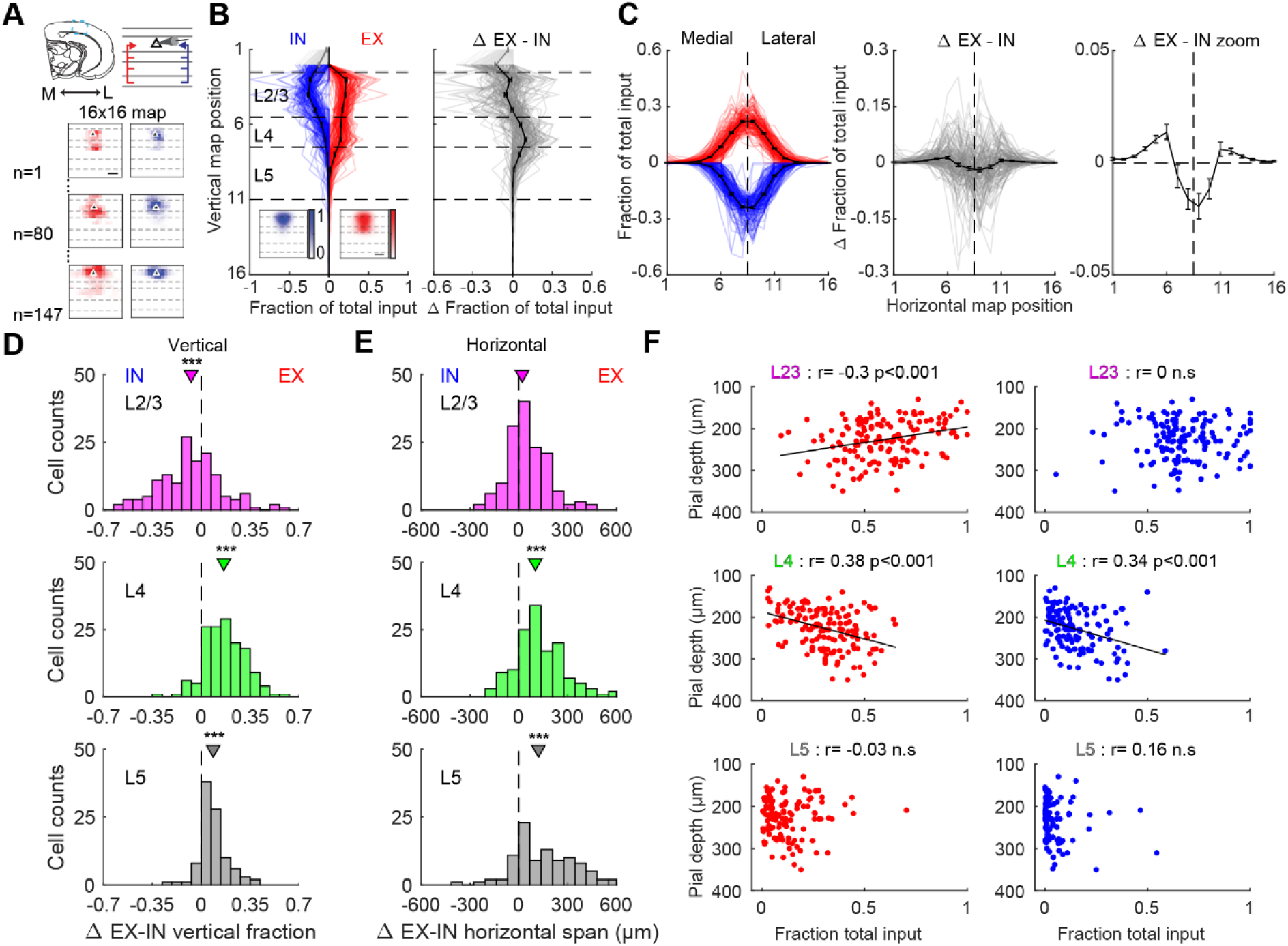
Spatial extent and overlap of local excitatory and inhibitory inputs to L2/3 pyramidal cells. **A** Alignment of input maps to the medial-lateral axis preceding analysis. Representative, peak normalized excitatory and inhibitory inputs maps for three cells (scale bar 100 µm). **B** Average vertical excitatory (EX, red) and inhibitory (IN, blue) input fraction per stimulus row; thin lines, individual cells (mean ± SEM, n= 147, left). Insets depict averaged normalized maps over all cells. Difference between vertical excitatory and inhibitory input fractions (right). Light grey areas in L1 not considered for comparison. **C** Same as B for horizontal excitatory and inhibitory input fraction per column and zoom-in of difference (right). **D** Distributions of differences between excitatory and inhibitory input fraction for L2/3, L4 and L5 (n=147). Triangles indicate mean. **E** Distributions of differences between horizontal extent of input origin for L2/3, L4 and L5 (n=147). Triangles indicate mean. **F** Excitatory (left) and inhibitory (right) input fractions plotted against pial depth for inputs from L2/3, L4 and L5 (n=147). Pearson correlation coefficient r indicated at top of each plot.

Next, we wanted to better understand if and how the input depends on a cell’s depth, i.e. its distance from the pial surface. As reported for auditory cortex (Meng et al. 2017), we observed that the fraction of excitatory and inhibitory input from L4 is correlated with the distance between the cell and the pia (Fig. 2F, middle row, r=0.38 and r=0.34, p<0.001, Pearson’s correlation coefficient) with more superficial cells receiving less fractional excitation and inhibition from L4 in comparison to deeper cells. In contrast, excitatory input from L2/3 displayed the opposite correlation (Fig. 2F, top row, r=-0.3, p<0.001, Pearson’s correlation coefficient). Such correlation was not present for L5 inputs and inhibitory input from L2/3 (L5 EX, r=-0.04, p=0.65; L5 IN, r=0.17, p=0.09; L2/3 IN, r=0, p=0.98, Pearson’s correlation coefficient). This indicates that upper L2/3 PCs, i.e. putative L2 cells, receive more excitatory input from L2/3 and less from L4 compared to lower layer L2/3 PCs, i.e. putative L3 cells that receive a larger fraction of input from L4.

### Parameters describing local inputs to L2/3 pyramidal cells

Sensory processing of visual stimuli, on a per-cell basis, is partially based on the integration of inputs that respond to stimuli at different locations in visual space, as observed for orientation tuning (e.g. Hubel and Wiesel 1962), and visual space is represented on the cortical sheet in a retinotopic fashion (Dräger, 1975; Wagor, Mangini, & Pearlman, 1980; Schuett, Bonhoeffer, & Hübener, 2002; Garrett et al., 2014). Selective sampling of visual space by individual neurons might therefore be reflected in the spatial organization of their inputs. Thus, we explored the vertical and horizontal spatial structure of the input maps in more detail. First, rather than considering the entire vertical and horizontal input fractions per layer, we determined the centroids of the input distributions and compared these for excitation and inhibition originating in the different layers (Fig. 3A-C).

**Figure 3:**
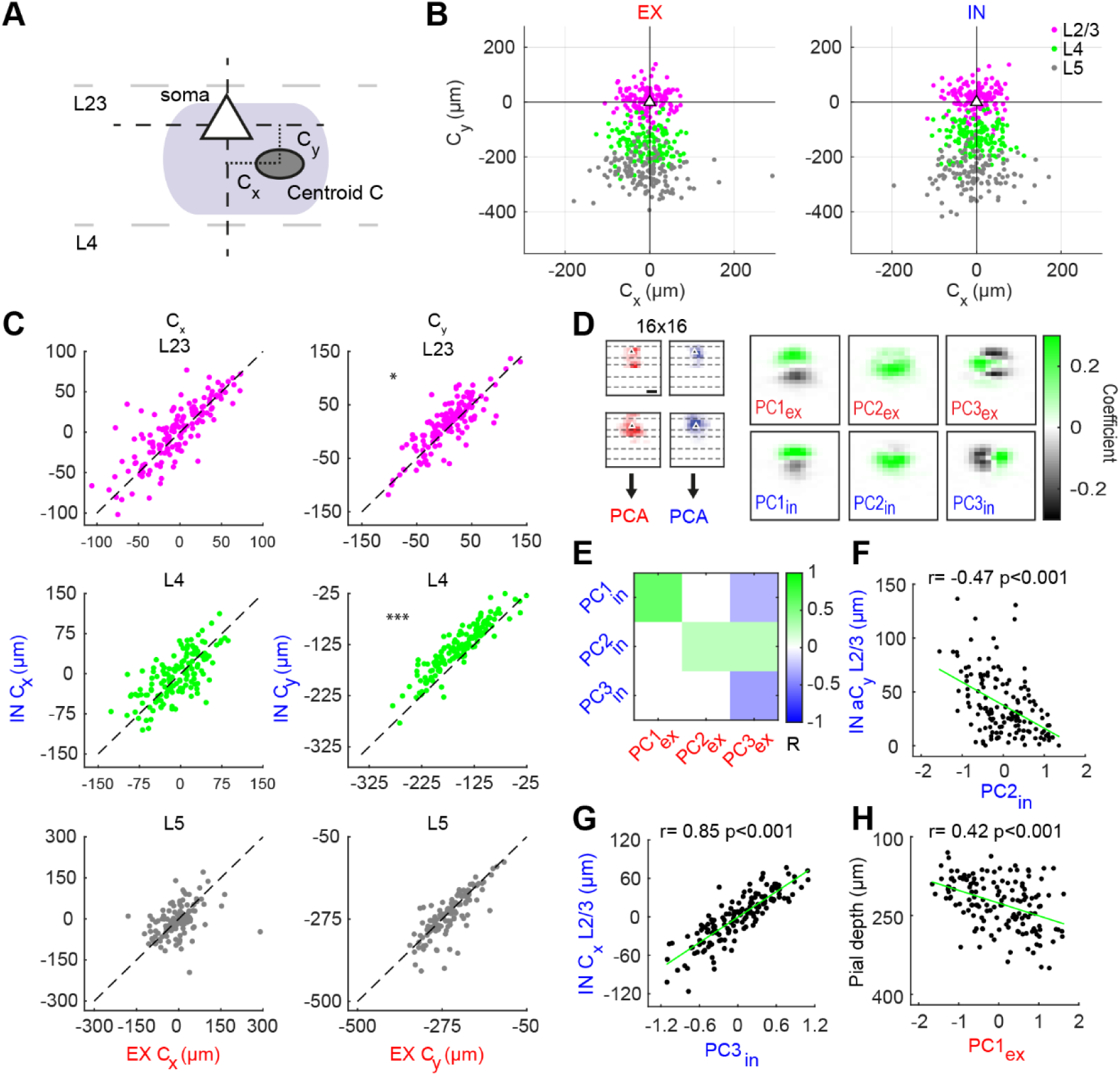
The relative position of the centroid explains most of the variance in the input maps. **A** Schematic depicting the position of the centroid (C) of an input map within L2/3 relative to the cell soma. The horizontal and vertical distances between soma and centroid were determined (C_x_ and C_y_). **B** C_x_ and C_y_ within L2/3, L4 and L5 for individual cells, both for excitation (left) and inhibition (right, n=147). **C** C_x_ (left column) and C_y_ (right column) of inhibition plotted against C_x_ and C_y_ of excitation, respectively, in L2/3, L4 and L5. Unity lines are indicated. Asterisks indicate significant differences. **D** Principal component analysis (PCA) using the 16×16 normalized excitatory and inhibitory input maps separately (left). Before PCA, the input maps were vertically and horizontally aligned (see Methods). Input maps corresponding to eigenvalues of the first three principal components for excitation and inhibition (right). **E** Correlations between the first three principal components for excitation and inhibition. Color indicates the Pearson correlation coefficient between the pair of parameters according to the color bar on the right. Coefficients with p values > 0.05 are set to 0. **F** Absolute C_y_ for inhibition plotted against PC2_in_ within L2/3 (n=147). **G** C_x_ for inhibition in L2/3 plotted against PC3_in_ (n=147)**. H** The distance of the cell from the pia plotted against PC1_ex_ (n=147).

The centroid was calculated as the arithmetic mean of the locations of the points in the input map, weighted by their input amplitude, to quantify the spatial offset of the synaptic input distribution in the different layers relative to the soma. The centroid position was described by its horizontal and vertical position with respect to the soma (C_x_ and C_y_, Fig. 3A). Figure 3B shows the centroid positions C_x_ and C_y_ for the excitatory (left) and inhibitory input (right) from L2/3, L4 and L5 for all cells relative to the soma. The centroid positions of excitation and inhibition showed similar distributions centered on the vertical axis. The horizontal spread of the distributions increased from L2/3 through L4 and to L5. C_x_ and C_y_ of neither excitatory nor inhibitory input centroids were correlated with each other within any layer (Supplementary Fig. 4A). While the excitatory and inhibitory C_y_ were significantly correlated with the distance between L2/3 somata and the pia across all layers, this relation was absent for C_x_ with the exception of the inhibitory input from L5 (Supplementary Fig. 4A). While on average the centroids of excitatory and inhibitory input followed each other in their position along the horizontal axis within all layers (Fig. 3C, left, Wilcoxon signed-rank, L2/3, p=0.38, n=147; L4, p=0.17, n=138; L5, p=0.24, n=98), there were a number of cells which showed horizontally displaced excitatory and inhibitory centroids. Vertically, the centroids of excitatory and inhibitory input were significantly different from each other for L2/3 and L4 but not for L5 (Fig. 3C, Wilcoxon signed-rank, L2/3, p=0.34, n=147; L4, p<0.001, n=138; L5, p<0.05, n=98). On average, the inhibitory centroids were above the corresponding excitatory centroids in L2/3 and L4 (Fig. 3C).

To gain further understanding of the information present in the input distributions of each cell, we applied principal component analysis (PCA) as an independent and unbiased method for identifying the primary spatial patterns that underlie the input maps. PCA was performed on the entire set of 16 x 16 pixel input maps, separately for excitation and inhibition (Fig. 3D). Before PCA, the input maps were horizontally and vertically aligned to the soma (see Methods). The input maps corresponding to the first three principal components (eigenmaps) for both excitatory and inhibitory inputs are displayed in Figure 3D. These three components together explained ∼60% variance in the data for both excitation and inhibition (Supplementary Fig. 4B). Inspection of these components revealed that the respective excitatory and inhibitory principal component weights were significantly correlated with each other, and in addition the third principal component of excitatory input PC3_ex_ was correlated with all inhibitory principal components (Fig. 3E). For comparison, we performed PCA on the combined excitatory and inhibitory maps, leading to similar results (Supplementary Fig. 4C-E). Moreover, the principal components were strongly related to the vertical and horizontal spatial features of the input maps already described above (Supplementary Fig. 4F). In general, these principal components described i) the relative laminar difference between input from upper and lower layers (PC1_ex_, PC1_in_), ii) the larger absolute fraction of input from lower layers (PC2_ex_, PC2_in_) and iii) the relative horizontal difference between input arising medial versus lateral from the soma (PC3_ex_, PC3_in_). For example, the PC2_in_ weights were significantly correlated with the vertical input offset described by the absolute distance between soma and inhibitory centroid (absolute value of C_y_, aC_y_, Fig. 3F, r=-0.4, p<0.001, Pearson’s correlation coefficient) and PC3_in_ was strongly correlated with the horizontal inhibitory input offset described by C_x_ (Fig 3G, r=0.81, p<0.001, Pearson’s correlation coefficient). Interestingly, four out of the six principal components were correlated with the pial depth, even though we accounted for the direct information about cell location before conducting PCA (Fig. 3H, PC1_ex_ vs. pial depth; r=0.42, p<0.001, Pearson’s correlation coefficient; Supplementary Fig. 4F). This indicates that the input pattern shape itself contains information about the cell location within L2/3.

In summary, the main factors characterizing both excitatory and inhibitory circuit motifs are the laminar differences between inputs as well as the vertical and horizontal input offsets described by C_y_ and C_x_.

### Relationship between visual response properties and input map structure

Next we explored whether the observed input circuit motifs were related to the visual response properties of individual neurons. Figure 4A shows the definition of stimulus orientation angles (0°, horizontal bars moving vertically; increasing angles follow clockwise rotation of the bar pattern orientation), as well as examples of both, eye-specific orientation tuning curves, and excitatory and inhibitory input maps for two selected cells. We acquired *in vivo* stimulus response properties and input maps from 70 cells, from which 54 were responsive to grating stimuli. Figure 4B shows a comparison of the distributions of the global orientation selectivity index (gOSI, see methods) and preferred orientation for all cells recorded *in vivo*, and for the subset for which input maps were acquired. Comparisons between other visual response parameters (ocular dominance, direction selectivity, preferred direction, tuning width and peak Ca^2+^ amplitude), as well as spontaneous activity parameters, are shown in Supplementary Figure 5.

**Figure 4:**
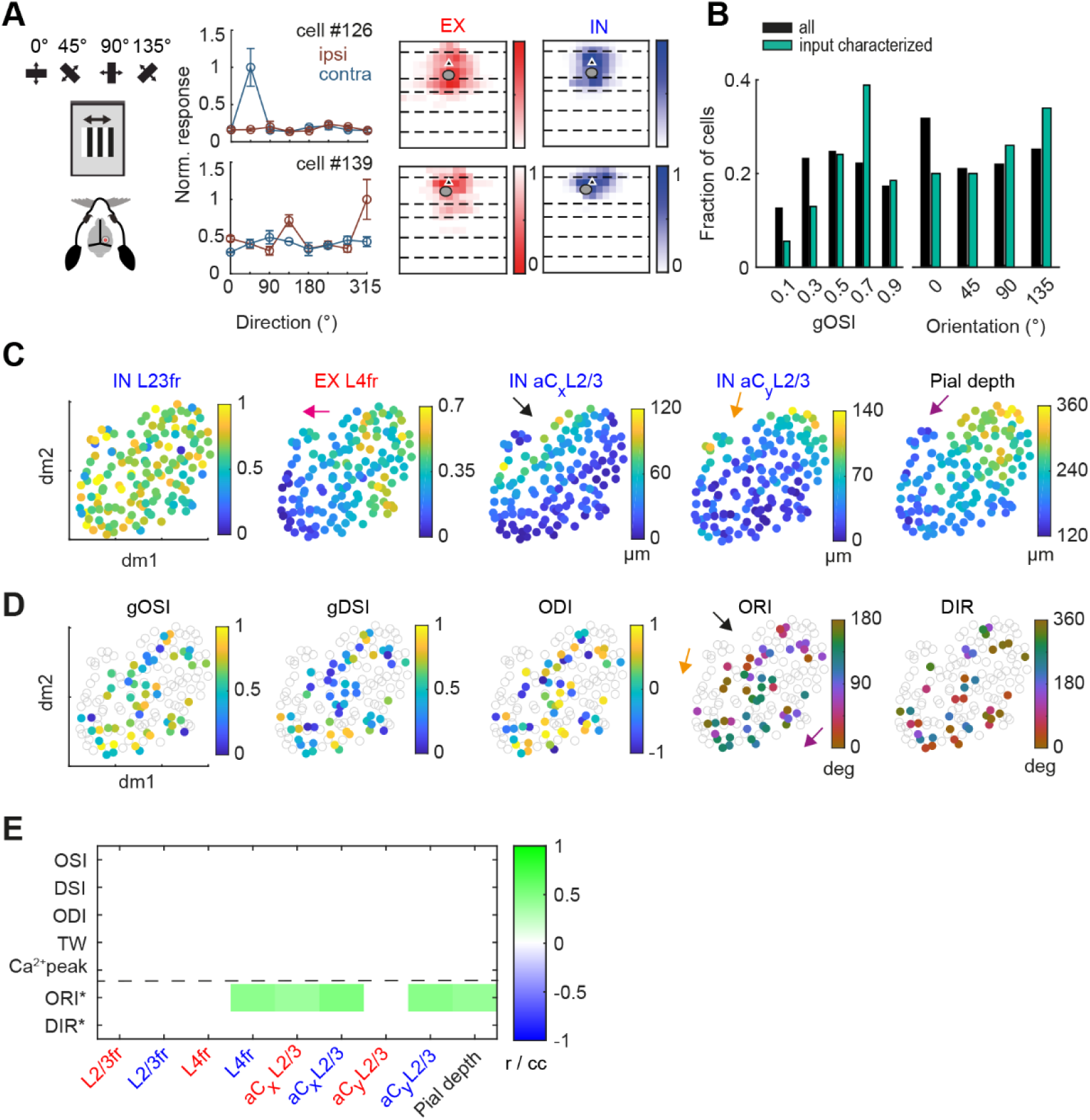
Relation between visual response properties and input map features. **A** Schematic of actual orientation of the moving grating stimuli with respect to the mouse and the recorded right hemisphere. 0° are horizontal gratings, increasing angles follow clockwise rotation of the stripe pattern orientation. Tuning curves for contra- (light blue) and ipsilateral (orange) eye stimulation of two example L2/3 PCs (mean ±SEM, n=4). Corresponding excitatory and inhibitory input maps (right). Input map centroids in L2/3 are indicated with grey circles. **B** Global orientation selectivity index (gOSI) and orientation preference (ORI) distributions of all *in vivo* characterized cells (black, n=1134, n=937) and subset of cells that were *in vivo* / *in vitro* characterized (turquoise, n=54, n=50). For the ORI distributions, only cells with gOSI>0.25 are included. **C** UMAP projections of different input map features (inhibitory L2/3 fraction, excitatory L4 fraction, absolute inhibitory C_x_ and C_y_ in L2/3 and pial depth, n=147). The UMAP embedding was performed using the excitatory L4 fraction, absolute inhibitory C_x_ as well as C_y_ in L2/3 and pial depth. Arrows indicate direction of gradients (dimensions 1 (dm1) and 2 (dm2) are plotted). **D** UMAP projections for gOSI, global direction selectivity (gDSI), ocular dominance index (ODI, n=54), ORI (n=50) and direction preference (DIR, n=38). For ORI and DIR, cells were subsampled based on gOSI>0.25 and gDSI>0.25, respectively. The embedding was performed using the same parameters used in C. **E** Correlations between visual response properties and input map features (TW: tuning width). Colors indicate the Pearson correlation or circular correlation coefficient (cc) between the pair of parameters according to the color bar on the right. Coefficients with p values > 0.05 are set to 0. For ORI and DIR, cells were subsampled based on gOSI>0.25 and gDSI>0.25, respectively.

For exploring the relationship between visual response properties and input organization, we displayed single cell data in two-dimensional UMAP (Uniform Manifold Approximation and Projection) plots, with an embedding based on a selection of the most distinctive parameters we had identified previously in the synaptic input maps (excitatory fraction from L4, inhibitory L2/3 aC_x_ and aC_y_, cf. Fig. 2-3), as well as pial depth (Fig. 4C, D). By color-coding the parameter values in the UMAP plots, we found specific gradients and groupings between the embedded input map parameters: The inhibitory L2/3 aC_x_ displayed a gradient which was mostly orthogonal to the gradients for inhibitory L2/3 aC_y_ and pial depth, illustrated by arrows in Figure 4C. The excitatory L4 fraction displayed a clear gradient which mostly followed the pial depth gradient. Individual parameters that were not used for embedding either followed the observed gradients, such as the excitatory aC_x_ within L2/3 and inhibitory aC_x_ within L4 (Supplementary Fig. 6A), or did not display any specific pattern, such as the inhibitory L2/3 fraction (Fig. 4C). Similar observations were made for the UMAP plots of the principal components where PC1-3_in_ as well as PC3_ex_ followed the observed gradients (Supplementary Fig. 6B), reflecting the strong correlations shown in Fig. 3F-H. When inspecting the visual response properties on the UMAP plot (using the same embedding based on the input map features), we found that of all visual response parameters, orientation preference (ORI) prominently followed multiple of the gradients described above, suggesting dependencies between input map features and orientation preference (Fig. 4C, Supplementary Fig. 6C). The apparent parameter dependencies are quantitatively assessed below (Fig. 4E). We found significant correlations between the aC_x_ and aC_y_ within L2/3 and orientation preference. In contrast, no other visual response property, such as orientation and direction selectivity, as well as ocular dominance, displayed a significant correlation (Fig. 4E, inhibition aC_x_ vs. ORI, cc=0.5, aC_y_ vs. ORI, cc=0.46, p<0.01, circular correlation).

In summary, preferred orientation was the strongest link we found between the spatial organization of input maps and functional response properties; in particular there was a positive correlation between orientation preference, aC_x_ and aC_y_. Additionally, the orientation preference displays a dependence on pial depth and on the input fraction from L4. Thus, circuit organization principles related to visual response properties are likely driven by multiple parameters acting in parallel.

### Relationship between spatial input map offset and orientation preference

To further explore the specific relation between preferred orientation and the spatial organization of synaptic inputs across layers (Fig. 5A), we focused on the centroid distribution of excitation and inhibition with respect to orientation preference (Fig. 5B). We observed that for cells preferring orientations around 125° the L2/3 centroids of excitatory and inhibitory input were located closer to the soma, and, as the radial distance from the soma increased, the orientation preference shifted to around 35° (Fig. 5B). This was also true for the inhibitory input within L4 (Fig. 5B, right).

**Figure 5:**
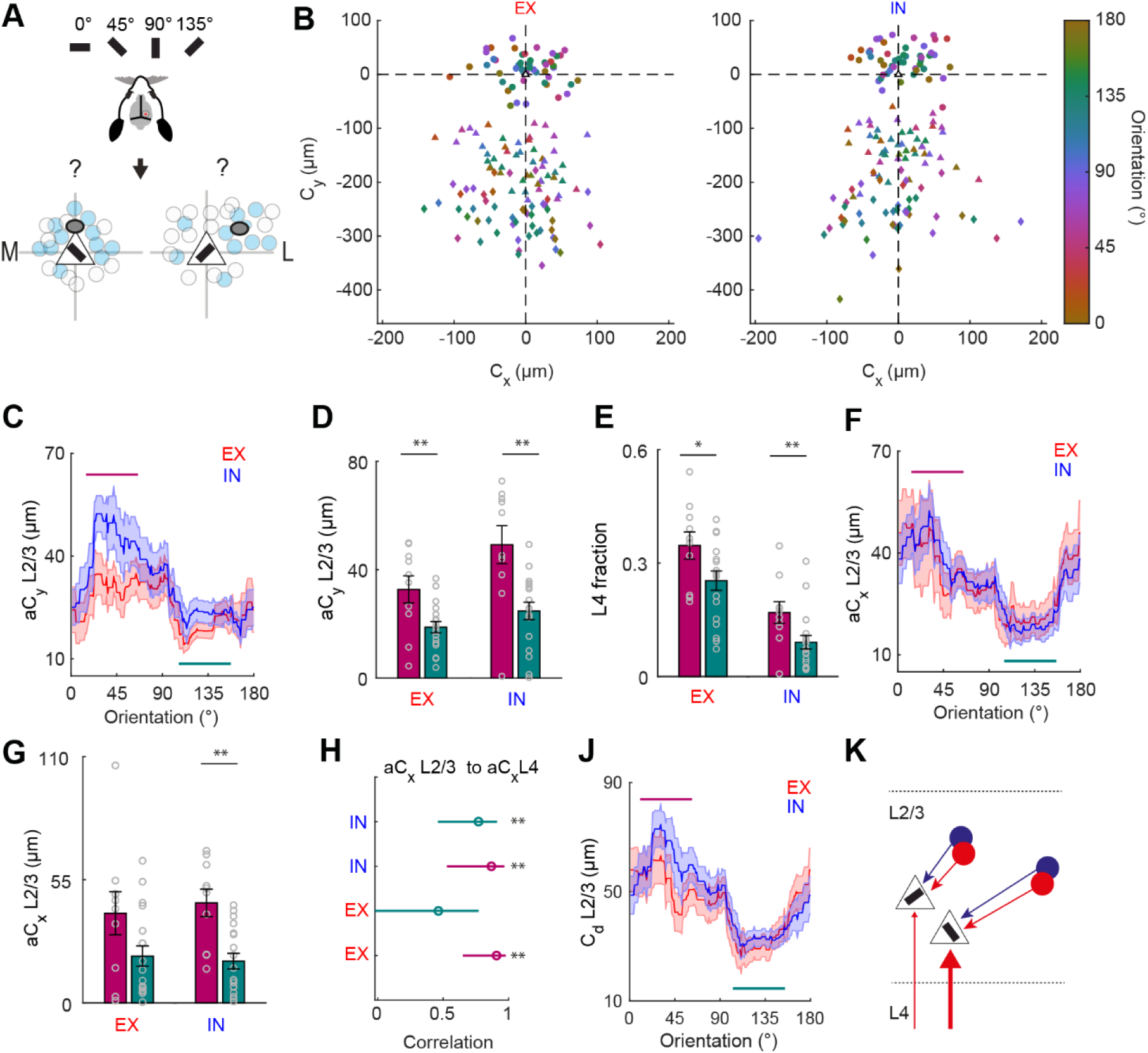
The synaptic input offset is linked to orientation preference. **A** Schematic displaying the gratings presented to the mouse, and the potential relative positions of the presynaptic input map centroid for two L2/3 PCs with orthogonal preferred orientations. **B** Centroid offsets C_x_ and C_y_ for L2/3 (circles), L4 (triangles) and L5 (diamonds) cells, color coded based on the cell’s preferred orientation (n=50) for excitation (left) and inhibition (right). Only cells with gOSI>0.25 are included. The white triangle marks the position of the cell soma. **C** Rolling average of excitatory (red) and inhibitory (blue) aC_y_ in L2/3 across orientation preference using a window size of 45°. The SEM is indicated as the shaded area. Turquoise and purple lines mark the angle ranges utilized for the comparison in the following panels (50° ranges: 10-60°, 100-150°, EX: cc=0.3, p=0.06, IN: cc=0.46, p<0.01). **D** Comparison of the average aC_y_ in L2/3 for the two highlighted sectors in panel C for excitation (left) and inhibition (right). Individual data points are superimposed (n=10, n=18). One-tailed Wilcoxon rank-sum. **E** Comparison of the average fraction of input originating in L4 for the two groups of angles indicated in panel C. One-tailed Wilcoxon rank-sum. **F** Same as in C for aC_x_ (n=50, EX: cc=0.39 p<0.05, IN: cc=0.5, p<0.001). **G** Same as in D for C_x_ (n=10, n=18). One-tailed Wilcoxon rank-sum. **H** Interlaminar (L2/3-L4) Pearson correlation of aC_x_ for excitation and inhibition. Error bars are 95% confidence intervals. **J** Same as in C for the distance between the soma of the cell and the centroid (C_d_, n=50, EX: cc=0.44 p<0.01, IN: cc=0.56, p<0.001). **K** Schematic summarizing observed vertical and horizontal input offsets for excitation and inhibition and its relation to orientation preference.

We first analyzed the relationship between the vertical component of the centroid offset and the preferred orientation, given the strong pial dependence we had found among input distribution parameters (Supplementary Fig. 4) and the potential presence of sublaminar subdivisions as observed in auditory cortex (Meng et al. 2017). For this, we computed the rolling average of aC_y_ across preferred orientation for excitatory and inhibitory maps within L2/3 (Fig. 5C). This distance displayed a sinusoidal relation to the orientation preferences (Fig. 5C). When comparing the two orientation ranges best separated by aC_y_, (∼35° and ∼125°), L2/3 PCs that preferred orientations ∼35° had a significantly larger vertical offset of their presynaptic inputs compared to cells preferring orientations ∼125°, for both excitation and inhibition (Wilcoxon rank-sum, p<0.01; Fig. 5D). Interestingly, the vertical centroid offset was more pronounced for inhibition compared to excitation (as observed above across the entire population, see Fig. 3C). Moreover, cells preferring ∼35° received stronger L4 synaptic input compared to cells preferring ∼125°, further distinguishing the input connectivity between these two groups of cells (Wilcoxon rank-sum, excitation: p<0.05, inhibition: p<0.01, Fig. 5E). To better understand this relationship, we computed rolling averages with the components of aC_y_, namely pial depth of the soma and depth of the centroid. Both of them were different between cells preferring ∼35°, located closer to the L4 border, and cells preferring ∼125°, located closer to the L1 border (Supplementary Fig. 7A-C), although pial depth correlated better with the separation. Therefore, while the best measure linking orientation preference and input map distribution vertically is aC_y_, this seems to be mostly determined by pial depth, with only a small component dependent on the relative positions of centroid and soma (Fig. 5K).

Next, we focused on the horizontal component of the centroid offset. PCs preferring ∼35° had a larger horizontal inhibitory input offset than cells preferring ∼125° (Wilcoxon rank-sum, excitation: p=0.06, inhibition: p<0.01, Fig. 5F, G). The horizontal inhibitory offset was also preserved across layers, with the horizontal position for the inhibitory centroids within L2/3 and L4 being significantly correlated for both groups of preferred orientations (Fig. 5B, H). In contrast to the vertical offset, the horizontal offset was independent of the soma location with respect to the pial surface (Supplementary Fig. 7D). As a last step, we quantified the overall synaptic input offset as the centroid distance to the soma (C_d_), and found an even stronger correlation to the preferred orientation (EX: cc=0.44 p<0.01, IN: cc=0.56, p<0.001, circular correlation) arguing that both the described vertical and horizontal input offset are linked to the overall orientation preference (Fig. 5J).

Taken together, these intracortical vertical and horizontal input offsets with respect to the postsynaptic cell (Fig. 5K) represent the first reported link between the functional intra- and interlaminar input distribution of a cell and one of its visual response properties, namely orientation preference.

### Relationship of orientation selectivity to dendritic morphology

Finally, we explored how the dendritic structure of L2/3 PCs was related to their tuning properties and intracortical connectivity. To this end, we reconstructed the dendritic morphologies of 97 L2/3 PCs for which we had determined their synaptic input connectivity. From 32 of these cells, we additionally obtained visual response properties.

PCs displayed a morphological continuum across L2/3 when considering the appearance of their apical tree (Supplementary Fig, 8A). PCs in lower L2/3 displayed a long apical dendrite with a tuft, whereas PCs in upper L2/3 had shorter but wider apical trees that branched profusely in L1 as previously described (Gouwens et al. 2019).

The apical and basal dendritic trees of L2/3 PCs have been shown to play distinct roles in sensory processing. Feed-forward inputs from L2/3 and L4 influence orientation preference of L2/3 PCs (Ko et al. 2011; Lee et al. 2016), most likely targeting basal dendrites (Young et al. 2019; Feldmeyer, Lübke, and Sakmann 2006). In contrast, cortico-cortical feedback inputs (Nassi, Lomber, and Born 2013; Wang et al. 2007; Smith et al. 2013) as well as orientation-tuned thalamocortical inputs (Chen et al. 2013; Roth et al. 2016) are likely to shape orientation selectivity via the apical dendrite. Therefore, we explored the relationship of apical and basal tree structure to tuning properties. The global orientation selectivity, tuning width as well as orientation preference of reconstructed L2/3 PCs covered the full parameter range displayed by all *in vivo* sampled cells (gOSI: 0.11-0.98, TW: 10.75-36.8°, ORI: 11.24-178.1°). Comparison of the examples of reconstructed dendritic morphologies of L2/3 PCs and their corresponding tuning curves in Figure 6A, B suggested that the apical tree morphology varies with tuning width.

**Figure 6:**
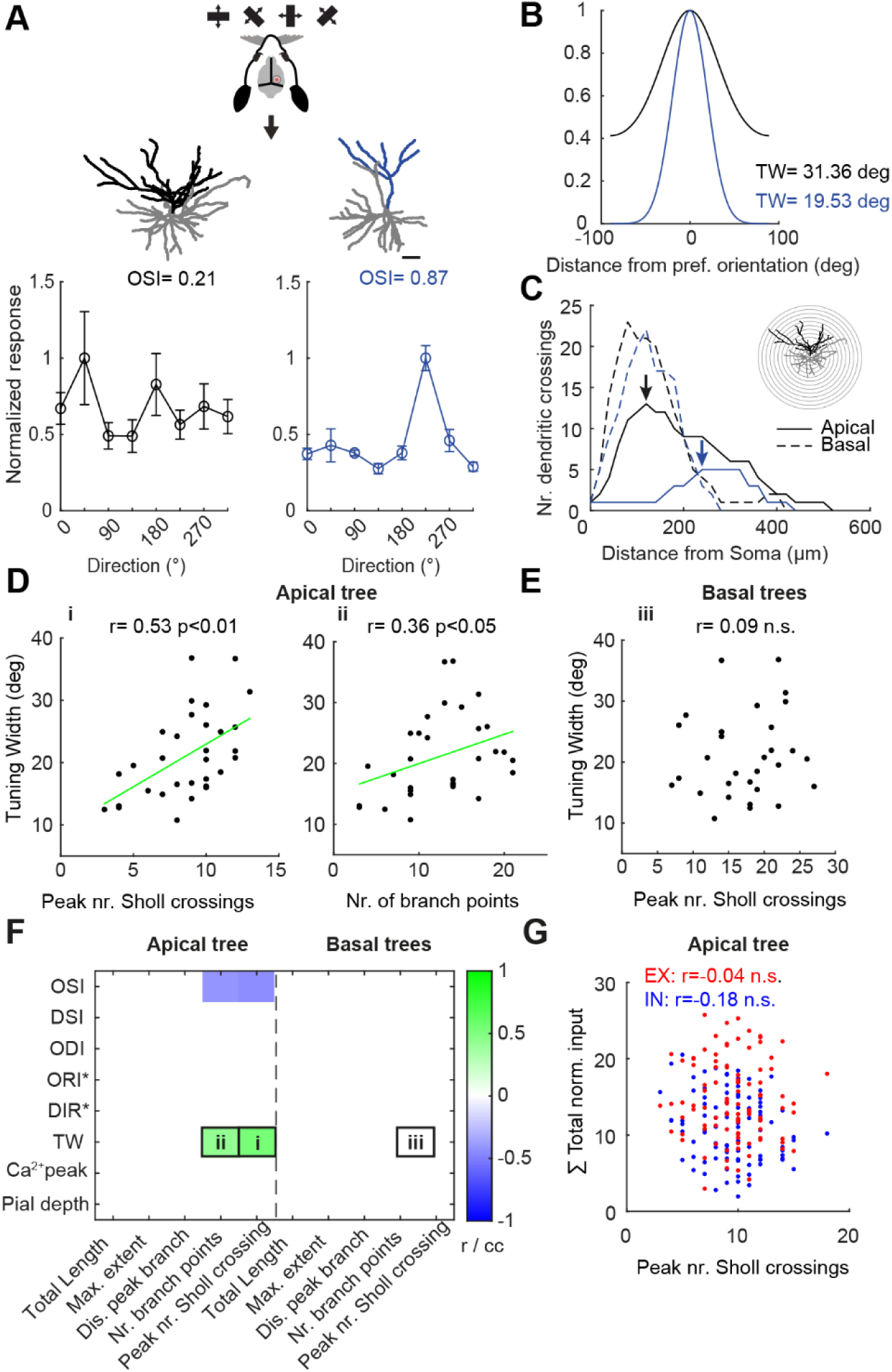
Apical but not basal dendritic complexity is related to the tuning width of L2/3 pyramidal cells. **A** Representative basal (grey) and apical dendritic morphologies (black/blue) of L2/3 PCs with low (left) and high orientation selectivity (right, pial depth: 190 and 220 µm). Normalized orientation tuning curves and OSI for the two depicted cells displayed at the bottom (mean ± SEM, n=4). **B** Gaussian fit of the tuning curves centered on the preferred orientation for the cells shown in A. Tuning width (TW) was determined as the full width at half maximum and is indicated on the right. **C** Sholl analysis for apical and basal dendritic trees for the two cells shown in A. The number of crossings was determined using concentric spheres centered around the soma at 20 µm increments. Arrows indicate the peak number of crossings of the apical trees. **D** TW plotted against the peak number of Sholl crossings (i in F) and against the number of branch points (ii in F) of the apical tree (n=32). **E** Tuning width plotted against the peak number of Sholl crossings of the basal trees (iii in F). **F** Correlations between visual response properties and morphological features. Colors indicate the Pearson correlation or circular correlation coefficient (cc) between the pair of parameters according to the color bar on the right. Total length, maximal extent and distance of peak branch (Dis. peak branch) are in µm. Coefficients with p values > 0.05 are set to 0. For ORI and DIR, cells were subsampled based on gOSI>0.25 and gDSI>0.25, respectively. **G** The sum of the total normalized excitatory input plotted against the peak number of Sholl crossings of the apical trees (n=97).

For a quantitative assessment of the spatial extent of the apical and basal tree, we extracted three morphological features related to dendritic length and two parameters related to dendritic complexity (Fig. 6C, Supplementary Fig. 8B, C, Sholl analysis, see Methods). Regarding dendritic complexity, we found a significant correlation between the tuning width and the peak number of Sholl crossings, as well as the total number of branch points of the apical tree (Fig. 6D, F, peak number of Sholl crossings, r=-0.53, p<0.01; number of branch points, r=0.36, p<0.05). Similar observations were made for the gOSI (Supplementary Fig. 8D, Fig. 6F). In contrast, the basal tree morphology did not show such correlations (Fig. 6E, F, Supplementary Fig. 8E, r=-0.09, p=0.52; r=0.15, p=0.53). Furthermore, for both the apical and basal trees, none of the measures for spatial extent or length significantly correlated with orientation selectivity nor tuning width (Fig. 6F). Moreover, none of the morphological parameters displayed a relation with the preferred orientation or other visual response properties (Fig. 6F). Noteworthy, while the maximal radial distance from the soma and the ratio between maximum horizontal and vertical extent of the dendrites displayed a pial depth dependency, the dendritic complexity as well as total length were independent from the pial depth of the soma (Fig. 6F). Finally, there was no relation between the summed excitatory or inhibitory input and apical as well as basal tree complexity (Fig. 6G, Supplementary Fig. 8F, apical, EX: r=-0.04, p=0.73; IN: r=-0,18, p=0.08, basal, EX: r=0.12, p=0.23, IN: r=-0,09, p=0.38).

Together, these results suggest that L2/3 PCs with narrow tuning width have a less complex apical dendritic tree than more broadly tuned L2/3 PCs. The complexity of the basal tree does not follow this rule.

## Discussion

We used a combined structure-connectivity-function analysis to directly link dendritic morphology and functional excitatory and inhibitory input patterns to the tuning properties of individual L2/3 PCs in binocular V1. We found that the strongest excitatory and inhibitory inputs to L2/3 PCs are within L2/3, while inputs from L4 and L5 are mainly dominated by excitation. Horizontally, the excitatory and inhibitory inputs follow the shape of an inverted Mexican hat, with inhibitory input originating from locations closer to the soma and excitatory input from more distant locations. Using unbiased feature extraction, we found that vertical and horizontal input offsets of excitation and inhibition are the features underlying most variance in the input distribution across cells. We quantified these spatial offsets via the input centroids. Exploring their relationship to visual response properties, we observed that centroid location was directly connected to the preferred orientation of the cells: L2/3 PCs with the largest difference between centroid offsets with respect to the soma preferred orientations that were orthogonal to each other. Moreover, the amount of L4 input to L2/3 PCs was different for cells preferring orthogonal orientations. This suggests that the strength as well as the spatial location of presynaptic inputs with respect to the postsynaptic soma are an important contributor to orientation preference. With respect to stimulus tuning properties, we found that orientation selectivity was directly related to the dendritic complexity of L2/3 PCs: Sharply tuned L2/3 PCs had a less complex apical tree compared to broadly tuned cells. Neither dendritic structure nor input connectivity was directly related to the ocular dominance of L2/3 PCs, arguing against eye-specific intracortical connectivity motifs at the level of L2/3.

Taken together, these results suggest an important role for both the intra- and interlaminar connectivity and dendritic structure of L2/3 PCs in shaping orientation tuning.

### Excitatory and inhibitory inputs to L2/3 PCs

A previous study on functional connectivity using the same LSPS approach for input mapping, suggested global spatial balance between excitation and inhibition across L2/3, L4 and L5 in V1 (Xu et al. 2016). In this study, however, the excitatory and inhibitory inputs were mapped in separate sets of L2/3 PCs. In the present study, we obtained the excitatory and inhibitory input to L2/3 PCs in the same cells, enabling a direct comparison of spatial overlap between excitation and inhibition. We found that, on average, excitatory and inhibitory inputs do indeed spatially overlap across layers, but on a single cell level the input organization is much more diverse and comparable to the diverse distribution of excitatory and inhibitory synaptic inputs to L2/3 PCs seen in auditory cortex (Meng et al. 2017). L2/3 PCs received stronger local inhibitory connections within L2/3 compared to excitation, inputs from L4 were dominated by excitation, and L5 excitatory inputs were rarely spatially balanced by inhibition or vice versa. Vertically, the strongest connections to L2/3 PCs were within L2/3 for both excitation and inhibition. Previous studies analyzing the number of presynaptic cells connected to L2/3 PCs using monosynaptic rabies tracing reported the highest number of presynaptic excitatory cells in L4 rather than L2/3 (Wertz et al. 2015; Rossi, Harris, and Carandini 2019). This discrepancy could be explained by the differences in purely structural vs. functional synapses formed between pre- and postsynaptic cells. While monosynaptic rabies tracing reveals structural connectivity, LSPS reveals functionally connected cells. Indeed, paired recordings show that the connection probability as well as synapse strength is higher between L2/3->L2/3 compared to L4-> L2/3 cells within V1 (Morgenstern, Bourg, and Petreanu 2016).

Horizontally, synaptic input distribution is organized as an inverted Mexican hat, with inhibitory input located close to the vertical axis through the L2/3 PC soma, and excitatory input extending more distally when including inputs across all layers. In L2/3, however, the horizontal extent was on average similar for excitation and inhibition, in contrast to L4 and L5. This differs from anatomical results obtained using cell reconstructions and monosynaptic rabies tracing, where excitatory inputs have been shown to display broader distribution relative to inhibition (Binzegger, Douglas, and Martin 2007; Rossi, Harris, and Carandini 2019). Again, this could be explained by the structure-function discrepancy.

The excitatory and inhibitory inputs with respect to cells in upper and lower L2/3 cells can be distinguished based on their ascending input from L4. L2/3 PCs close to the L4 border received more L4 excitatory as well as inhibitory input compared to L2/3 PCs close to the L1 border. This is similar to the auditory cortex, where L2/3 PCs were subdivided into L2 and L3 based on their degree of L4 input (Meng et al. 2017) arguing for common circuits schemes across sensory brain areas.

### Relationship of intra- and interlaminar circuits to tuning properties of L2/3 PCs

How orientation tuning is computed or amplified through intracortical excitatory and inhibitory circuits in L2/3 is still debated. Several studies have demonstrated that local as well as long-range intracortical connectivity of similarly tuned cells follows certain rules. For example, L2/3 PCs responding to similar stimulus features share strong local excitatory reciprocal connections within V1 (Cossell et al. 2015). Moreover, besides the strength of connectivity, the location of presynaptic receptive fields in visual space has been shown to correlate with a cell’s orientation tuning: Neurons preferring the same orientation tend to connect to each other when their receptive fields are aligned along the axis of their preferred orientation (Schwarz and Bolz 1991; Bosking et al. 1997; Iacaruso, Gasler, and Hofer 2017; Rossi, Harris, and Carandini 2019). Our results demonstrate that the spatial arrangement of presynaptic soma locations in cortical space, weighted by the synaptic input strength, are related to the preferred orientation of the postsynaptic cell. We observe both a vertical and a horizontal input offset, which most likely can be attributed to different circuit mechanisms.

According to retinotopic mapping, a bar with an orientation of ∼135° would align along a coronal brain slice (as we have used), whereas a bar of the orthogonal orientation (∼45°) would be pointing into and out of the slice (Garrett et al. 2014). This arrangement, together with the aforementioned relationship between cortical space layout of the presynaptic cells and orientation preference of the postsynaptic cell, yields two hypotheses: 1) the horizontal extent of the presynaptic input should differ between the aligned and the orthogonal bar. 2) Any anisotropic distribution of presynaptic cells in cortical space related to the postsynaptic cell’s preferred orientation should become apparent. We found that the horizontal span of the spatial input distribution indeed tends to be larger for PCs preferring the orientation aligned with the slice (∼125°) compared to PCs preferring the orthogonal orientation (Supplementary Figure 6B). In addition, in PCs preferring the orthogonal orientation (∼35°) the input centroids are more offset from the soma (Figure 5B), meaning these input distributions have an asymmetry perpendicular to the long axis of the input distribution. Because this asymmetry would point into/out of the slice in PCs preferring the orientation aligned with the slice (∼135°), their apparent input centroids are located close to the soma in our preparation. However, how our measurements in cortical space compare to the detailed distribution of the presynaptic cells in visual space must be measured in future studies.

Besides horizontal connectivity, the vertical input within and across layers could potentially influence the orientation tuning. Generally, anisotropies in the preferred orientation of neurons have been observed within L2/3 and across other layers in visual cortex (Kreile, Bonhoeffer, and Hübener 2011; Sun et al. 2016). The bias in orientation preference of L2/3 PCs in the upper and lower part of L2/3 observed in our study (Supplementary Fig.7 A-C, E) was accompanied by a difference in vertical input connectivity from L2/3 and L4. Lower L2/3 PCs could in principle directly inherit their orientation preference from connected L4 cells. We found that L2/3 PCs that prefer orientations of ∼35° indeed receive stronger synaptic input from L4, and are also located mostly in the lower part of L2/3. Cells preferring the orthogonal orientation of ∼125° could be embedded in a different cortical network.

We found that, while both the excitatory and inhibitory inputs spatially overlap to a large degree, the inhibitory input offset is more strongly related to the orientation preference than the excitatory offset. This could indicate that orientation preference is linked to specific inhibitory connectivity. Inhibitory cells have been shown to strongly inhibit those pyramidal cells that provide them with strong excitation and share their response selectivity, indicating that there is specific inhibitory connectivity in a recurrent intracortical network (Znamenskiy et al. 2018; Cossell et al. 2015).

### Orientation selectivity and dendritic structure

We found that sharply orientation tuned L2/3 PCs have a less complex apical tree compared to broadly tuned cells. In general, it has been demonstrated that tuning width correlates with the degree of dendritic non-linearities caused by clustering of spines with similar tuning (Wilson et al. 2016; Smith et al. 2013). This applies to both the apical and basal tree. It is conceivable that such non-linearities become more important the fewer dendritic branches a L2/3 PC has. However, we only found a significant correlation between the apical, not basal, dendritic complexity and tuning width. Therefore, it is likely that additional mechanisms shaping orientation selectivity are specific to the apical dendrite. Whereas basal dendritic trees receive local feedforward input via L2/3 and L4, apical dendrites of L2/3 PCs in V1 receive orientation tuned input via at least four different sources: 1) projections from the LGN (Cruz-Martín et al. 2014) 2) projections from lateral posterior thalamus (Roth et al. 2016) 3) feedback cortico-cortical projections from higher visual areas (Nassi, Lomber, and Born 2013; Wang et al. 2007; Yang et al. 2013) and 4) inhibitory inputs targeting the apical dendrite (Chiu et al. 2013). These input sources carry diverse tuning, and have been shown to directly modulate the orientation selectivity of L2/3 PCs. Silencing of feedback projections from higher visual areas has been shown to reduce orientation selectivity, without altering the orientation preference of the postsynaptic cell (Nassi, Lomber, and Born 2013).

A plausible explanation for our observation is that a more complex apical dendrite enables denser sampling from nearby axons per area, given the larger axodendritic overlap (Shepherd et al. 2005). This would favor the formation of more synapses between presynaptic inputs with diverse tuning, originating from the above listed sources. Therefore, the integration of these diverse inputs would lead to a broader somatic tuning of the postsynaptic L2/3 PC. A less complex apical tree would have less axodendritic overlap, and simply sample from less nearby axons, leading to less modulatory effect on the postsynaptic orientation tuning from input sources 1-4 (Shepherd et al. 2005). Importantly, long-range feedback or feedforward input cannot be activated via glutamate uncaging. We do not find any relation between total synaptic input measured with LSPS and apical tree complexity in this study, supporting the idea of the modulatory effect of long-range inputs.

In conclusion, we show that specific visual response properties of L2/3 principal cells in mouse visual cortex are related to trans- and intralaminar circuit connectivity motifs as well as cellular morphology.

## Methods

### Animals

All experimental procedures were carried out in compliance with institutional guidelines of the Max Planck Society and the local government (Regierung von Oberbayern). Wild type C57bl/6 female mice (postnatal days P28-P70) were used. Craniotomy, virus injections and head plate implantation were performed at P30-P35. *In vivo* imaging and subsequent *in vitro* brain slice experiments were performed at P50-P70.

### Virus preparation and dilution

The GECI AAV2/1-Syn-FLEX-mRuby2-CSG-P2A-GCaMP6m-WPRE-SV40 (titer: 2.9 x 10^13^ GC per ml, Addgene accession no. 102816) in combination with the Cre recombinase AAV2/1.CamKII0.4.Cre.SV40 (titer: 1.8 x 10^13^ GC per ml, University of Pennsylvania Vector Core accession no. AV-1-PV2396) were used. The final titer of AAV2/1-Syn-FLEX-mRuby2-CSG-P2A-GCaMP6m-WPRE-SV40 was 1.4 x 10^13^ GC per ml (PBS was used for dilution).

### Solutions

Cortex buffer for *in vivo* surgeries and imaging contained 125 mM NaCl, 5 mM KCl, 10 mM glucose, 10 mM HEPES, 2 mM CaCl_2_ (2 ml 1M CaCl_2_) and 2 mM MgSO_4_ (2 ml 1M MgSO_4_). The buffer was sterilized and maintained at pH 7.4.

The cutting solution for *in vitro* experiments contained 85 mM NaCl, 75 mM sucrose, 2.5 KCl, 23 mM glucose, 1.25 mM NaH_2_PO_4_, 4 mM MgCl_2_, 0.5 mM CaCl_2_ and 24 mM NaHCO_3_, 310-325 mOsm, bubbled with 95% (vol/vol) O_2_, 5% (vol/vol) CO_2_. Artificial cerebrospinal fluid (ACSF) contained 127 mM NaCl, 2.5 mM KCl, 26 mM NaHCO_3_, 2 mM CaCl_2_, 2 mM MgCl_2_, 1.25 mM NaH_2_PO_4_ and 11 mM glucose, 305-315 mOsm, bubbled with 95% (vol/vol) O_2_, 5% (vol/vol) CO_2_. Caesium-based internal solution contained 122 mM CsMeSO_4_, 4 mM MgCl_2_, 10 mM HEPES, 4 mM Na-ATP, 0.4 mM Na-GTP, 3 mM Na-L-ascorbate, 10 mM Na-phosphocreatine, 0.2 mM EGTA, 5 mM QX-314, and 0.03 mM Alexa 594, pH 7.25, 295-300 mOsm. K-based intracellular recording solution contained 126 mM K-gluconate, 4 mM KCl, 10 mM HEPES, 4 mM Mg-ATP, 0.3 mM Na-GTP, 10 mM Na-phosphocreatine, 0.3-0.5% (wt/vol) Neurobiotin tracer and 0.03 mM Alexa 594, pH 7.25, 295-300 mOsm.

### Virus injection and chronic window preparation

The detailed procedure is described elsewhere (Weiler et al. 2018). Briefly, surgeries were performed on 31 female C57bl/6 mice (postnatal days P27-P35) that were intraperitoneally (i.p.) anesthetized with a mixture of Fentanyl (0.075 mg kg^-1^), Midazolam (7.5 mg kg^-1^) and Medetomidine (0.75 mg kg^-1^). Additional drugs applied were Carprofen (4mg/kg, subcutaneous, s.c.) before surgery and Lidocaine (10%, topical to skin prior to incision). A section of skin over the right hemisphere starting from the dorsal scalp was removed and the underlying periosteum tissue was carefully removed. A custom machined metal head bar (oval shape, with an 8 mm opening and two screw notches) was carefully placed and angled over the binocular zone of the primary visual area. The precise location of the binocular zone was determined by intrinsic optical signal (IOS) imaging through the intact skull prior to the craniotomy in each animal (see section below). A circular craniotomy (4 mm diameter) centered over the binocular zone of the right primary visual cortex was performed. The premixed virus was injected 200-500 µm below the pial surface at a single site in the binocular zone of V1 (50-100 nl/injection, ∼ 10 nl/min ejected by pressure pulses at 0.2 Hz, using glass pipettes and a pressure micro injection system. Additionally, diluted fluorescent retrobeads (1:20 with cortex buffer, Lumafluor Inc.) were pressure injected (10-20 nl/injection, 5 nl/min) medial and lateral to the virus injection site at ∼1500 µm from its center. The craniotomy was covered with a glass cover slip and was sealed flush with drops of histoacryl. The head bar and cover glass were then further stabilized by dental cement. After surgery, the animal was injected s.c. with saline (500 µl) and the anesthesia was antagonized by i.p. injection of Naloxone (1.2 mg kg^-1^), Flumazenil (0.5 mg kg^-1^) and Atipamezole (2.5 mg kg^-1^). Cells were allowed to express the virus for at least 2 weeks before *in vivo* imaging. Carprofen (4mg/kg, subcutaneous, s.c.) was administered the following two days.

### Intrinsic optical signal imaging

For IOS imaging, the optical axis was orthogonal to the head bar for each animal. The brain surface was first illuminated with light of 530 nm to visualize the blood vessel pattern and subsequently with 735 nm for intrinsic imaging in order to localize the BZ. Images were acquired using an x4 air objective (NA 0.28, Olympus) and a high-speed CCD camera (12 bit, 250×348 pixel, 40 Hz). The camera was focused ∼500 µm below the pial surface. Image acquisition and analysis software were custom written in Matlab. A patch with a size 20° x 40° was displayed randomly to either the left or the right mouse eye at two distinct positions next to each other in the central visual field. The patch was a sinusoidal grating displayed in eight directions for 7 s (grating orientation was changed every 0.6 s) with a temporal frequency of 2 cycles/s and a spatial frequency of 0.04 cycles/degree. A blank grey screen (50% contrast) was displayed for 5 s between each stimulus presentation. Individual trials were separated by 8 s and the entire stimulus sequence was repeated at least 2 times per eye and patch position during the surgery and at least 3 times per eye and patch position during the first *in vivo* imaging session

### *In vivo* 2-photon imaging

L2/3 PCs co-expressing GCaMP6m and a bright structural marker mRuby2 (mRuby2-CSG-P2A-GCaMP6m) were imaged *in vivo* using a tunable pulsed femtosecond Ti:Sapphire laser (Newport Spectra-Physics). The 2-photon laser was tuned to λ=940 nm in order to simultaneously excite GCaMP6m and mRuby2. An x16 0.8 NA water immersion objective was used to detect red and green signals. The excitation light was short passed filtered (720/25 short-pass) and the emitted photons passed through a primary beam splitter (FF560 dichroic) and green and red band pass filters onto GaAsP photomultiplier tubes.

Multiple imaging planes were acquired by rapidly moving the objective in the z-axis using a high-load piezo z-scanner. The image volume for functional cellular imaging was 250 x 250 x 100 µm^3^ with 4 inclined image planes that were each separated by 25 µm in depth. Imaging frames of 512 x 512 pixels (pixel size 0.49 µm) were acquired at 30 Hz by bidirectional scanning of an 8 kHz resonant scanner while beam turnarounds were blanked with an electro-optic modulator (Pockels cell). Imaging was performed between 130-400 µm below the pial surface. Excitation power was scaled exponentially (exponential length constant ∼150 µm) with depth to compensate for light scattering in tissue with increasing imaging depth. The average power for imaging was <50 mW, measured after the objective. The optical axis was adjusted orthogonal to the cranial window. ScanImage 4.2 (Pologruto, Sabatini, and Svoboda 2003) and custom written hardware drivers were used to control the 2PLSM microscope.

After functional characterization of L2/3 PCs, at least two high-resolution structural image stacks with different field of view sizes (low and high) were acquired at λ=940 nm/1040 nm. These stacks were acquired from the pial surface to L5/L6 and contained the functionally characterized L2/3 pyramidal cells of interest. These structural stacks usually consisted of 1) 450 sections (512 x 512 pixels) with a pixel size of 0.5 µm collected in z-steps of 1.4 µm (resulting in an imaged volume of 256 x 256 x 630 µm^3^). 2) 350 sections (512 x 512 pixels) with a pixel size of 1.9 µm collected in z-steps of 2 µm (resulting in an imaged volume of 972 x 972 x 700 µm^3^).

Experiments were performed under light anesthesia. Data acquisition started ∼45 min after mice were injected with an i.p. injection of Fentanyl (0.035 mg kg^-1^), Midazolam (3.5 mg kg^-1^) and Medetomidine (0.35 mg kg^-1^). Additional anesthetics (25% of induction level) were subcutaneously injected every 45-60 mins to maintain the level of anesthesia. Ophthalmic ointment was applied to protect the eyes. Mice were fixed under the microscope by screwing the metal head-plate to two posts and stable thermal homeostasis was guaranteed by using a heated blanket throughout the imaging session. Eye and pupil positions were recorded with two cameras throughout imaging.

### Visual stimulation

Visual stimuli were generated using the MATLAB Psychophysics Toolbox extension and displayed on a gamma-corrected LCD monitor (http://psychtoolbox.org). The screen measured 24.9 x 44.3 cm, had a refresh rate of 60 Hz and was positioned in portrait 13 cm in front of the eyes of the mouse. The monitor was adjusted in position (rotation and tilt) for each mouse to cover the binocular visual field. The presented stimuli area was chosen to subtend binocular visual space and the rest of the screen was uniformly grey (50% contrast). An OpenGL shader was applied to correct for the increasing eccentricity on a flat screen relative to the spherical mouse space (Marshel et al. 2011). Monocular stimulation of the eyes was achieved by servo-motor driven eye shutters that were operated by a microcontroller (see: http://csflab.nin.knaw.nl/protocols/eyeshutters) and MATLAB.

For all visual stimuli presented, the backlight of the LED screen was synchronized to the resonant scanner to turn on only during the bidirectional scan turnaround periods when imaging data were not recorded (Leinweber et al. 2014). The mean luminance with 16 kHz pulsed backlight was 0.01 cd/m^2^ for black and 4.1 cd/m^2^ for white.

To measure ocular dominance, the right or left eye was visually stimulated in random order using sinusoidal gratings of eight directions with a temporal frequency of 3 cycles/s and a spatial frequency of 0.04 cycles/degree. In order to cover the binocular visual space, the visual stimuli were presented at −25° to 25° azimuth and −15° to 35° elevation relative to the midline. Stimulation duration for moving gratings was 5 s interleaved by 6 s of a full-field grey screen. Trials were repeated 4 times per eye and direction.

Spontaneous activity was measured during 10 min in complete darkness with the monitor being turned off and eye shutters removed.

### Acute brain slice preparation and reidentification of cells

The detailed procedure is described elsewhere (Weiler et al. 2018b). Briefly, naïve mice (4-8 weeks old) and mice 1-2 days after *in vivo* imaging were deeply anesthetized with Isoflurane in a sealed container (>100 mg/kg) and rapidly decapitated. Coronal sections of V1 (320 µm, Bregma −1.5 to −3) were cut in ice cold carbogenated cutting solution using a vibratome (VT1200S, Leica). Slices were incubated in cutting solution in a submerged chamber at 34°C for at least 45 min and then transferred to ACSF in a light-shielded submerged chamber at room temperature (21°C) until used for recordings. Brain slices were used for up to 6 hours. A single brain slice was mounted on a *poly-D-lysine coated* coverslip and then transferred to the recording chamber of the *in vitro* 2PLSM while keeping track of the rostro-caudal orientation of the slice. For *in vivo* / *in vitro* experiments, the fluorescence bead deposits in the brain slice where used to locate the area of interest by comparing the recorded distance between beads and imaging area to the ones obtained *in vivo*. Following this, a high-resolution image stack was acquired from the slice surface to the bottom using an x16 objective and a wavelength of 1040 nm to excite mRuby2. ScanImage 4.2 and custom written hardware drivers were used to operate the *in vitro* 2PLSM microscope. The *in vitro* stack consisted of 200-320 sections (512 x 512 pixels; 0.5 −2 µm per pixel) recorded in z steps of 1-2 µm. As a next step, the relative positions of cells and morphological details such as blood vessel patterns were compared between the side view of the *in vivo* stack and the face view of the *in vitro* stack. Z-projections of sections of the *in vivo* side view and *in vitro* stacks were created (50 sections with 1 µm spacing using Image J) and used to compare and match cell patterns in z-projections by eye.

### Photostimulation

For uncaging experiments using UV laser light two different setups were used. Coronal brain slices were visualized with an upright microscope (setup 1: BW51X, Olympus, setup 2: A-scope, Thorlabs) using infrared Dodt gradient contrast (DGC) with a low magnification UV transmissive objective (4x objective lens) and images were acquired by a high-resolution digital CCD camera. The digitized images from the camera were used for registering photostimulation sites in cortical brain sections. MNI-caged-L-glutamate concentration was 0.2 mM. The bath solution was replaced after 3 h of recording, and bath evaporation was counterbalanced by adding a constant small amount of distilled H_2_O to the solution reservoir using a perfusor. For *in vitro* experiments without previous cell characterization *in vivo*, L2/3 PCs in bV1 were primarily targeted using morphological landmarks and then whole cell recordings were performed at high magnification using a 60x objective. Targeted PC bodies were at least 50 µm below the slice surface. For the *in vivo* / *in vitro* experiments, 2-photon guided targeted patching was performed on cells that were matched *in vivo* and *in vitro*. Borosilicate glass patch pipettes (resistance of 4-5 MΩ) were filled with a Cs-based internal solution for measuring excitatory and inhibitory postsynaptic currents (EPSC: voltage clamped at −70 mV, IPSC: voltage clamped at 0-5 mV). Electrodes also contained 30 µM Alexa 594 for detailed morphological visualization using 2-photon microscopy. Once stable whole-cell recordings were obtained with good access resistance (< 30 MΩ) the microscope objective was switched from 60x to 4x. For circuit mapping, the slice was positioned within the CCD camera’s field of view and a stimulus grid (16 x 16 with 69 µm spacing) was aligned to the pial surface using Ephus software (Suter et al. 2010). The location of the cell soma was noted in Ephus. The UV laser power was adjusted to 10-15 mW in the specimen plane and then the mapping was initiated (1 ms pulses, 1s interstimulus interval). Multiple maps were recorded in a pseudo-random fashion while clamping the cell at −70 mV (2-3 repetitions with change of mapping sequence during each trial). Optionally, multiple (2-3 repetitions) inhibitory laminar input maps were recorded at 0 mV.

On setup A (SA), a diode pumped solid state (DPSS laser Inc.) laser was used to generate 355 nm UV laser pulses for glutamate uncaging. The duration and intensity of the laser pulses were controlled by an electro-optical modulator, a neutral density filter wheel and a mechanical shutter. The beam of light was controlled using voltage-controlled mirror galvanometers. An UV-sensitive photodiode measured the power of the UV laser beam. A dichroic mirror reflected the UV beam into the optical axis of the microscope while transmitting visible light for capturing bright-field images by the CCD camera. The beam passed a tube/scan lens pair in order to underfill the back aperture of the x4 mapping objective resulting in a pencil-shaped beam.

On setup B (SB), the UV laser for glutamate uncaging was an Explorer One 355-1 (Newport Spectra-Physics). The duration and intensity of the laser pulses were directly controlled using analog signals and the built-in software L-Win and a mechanical shutter as well as neutral density filters. An UV-sensitive photodiode measured the power of the UV laser beam.

Data were acquired with Multiclamp 700 B amplifiers (Axon instruments). Voltage clamp recordings were filtered at 4-8 khz and digitized at 10 kHz. Data Analysis was performed using custom-written software in MATLAB. The spatial resolution of photostimulation was estimated using excitation profiles (Shepherd and Svoboda 2005). Excitation profiles describe the spatial resolution of uncaging sites that generate action potentials in stimulated neurons. For this, excitatory as well as inhibitory cells in different layers of V1 were recorded either in whole-cell or cell-attached configuration using the K-based internal solution with the amplifier in current-clamp mode. The microscope objective was then switched from 60x to 4x and a 8×8 or 8×16 stimulus grid with 50 or 69 µm spacing was overlaid on the slice image and the soma location was registered. The interstimulus interval was set to 1 s and a map was acquired.

### Image acquisition for morphological imaging

The patch pipette was carefully retracted from the cell after successful recording and filling with Alexa-594. A detailed structural 2-photon image stack of the dendritic morphology of the entire cell was acquired with excitation light of λ=810 nm using ScanImage 4.2 (Pologruto, Sabatini, and Svoboda 2003). The structural image stacks typically consisted of 250 sections (1024 x 1024 pixels; 0.3-0.8 µm per pixel) collected in z steps of 1-2 µm. For cells that contained mRuby2 as structural marker, a second identical image stack was acquired at λ=940/1040 nm. An overlay of the acquired stacks (in ImageJ) was then used to verify that the *in vivo* functionally characterized cell of interest was successfully re-identified, recorded and filled with Alexa 594.

### *In vivo* imaging analysis

Custom-written Matlab software was used for image and data analysis. For optical signal imaging analysis, the acquired images were high-pass filtered and clipped (1.5%) to calculate blank-corrected image averages for each condition. Additionally, a threshold criterion (image background mean + 4 x standard deviation) was set to determine the responsive region within the averaged image. The mean background value of the non-responsive region was subtracted from each pixel and all pixel values within the responsive area were summed to obtain an integrated measure of response strength.

The use of GCaMP6m in combination with mRuby2 gave the possibility to perform ratiometric imaging. Image sequences were full-frame corrected for tangential drift and small movements caused by heart beat and breathing. An average of 160 image frames acquired without laser excitation was subtracted from all frames of the individual recording to correct for PMT dark current as well as residual light from the stimulus screen. Cell body detection was based on the average morphological image derived from the structural channel (mRuby2) for each recording session. ROIs (Region of interest) were drawn manually, annotated and re-identified in subsequent imaging sessions. The fluorescence time course of the area within the cell body was calculated by averaging all pixel values with the ROI on both background-corrected channels. Cell calcium traces were then low-pass filtered (0.8 Hz cut-off) and the neuropil signal subtracted using a neuropil factor r of 0.7(Kerlin et al. 2010). The green and red fluorescence signal were estimated as:

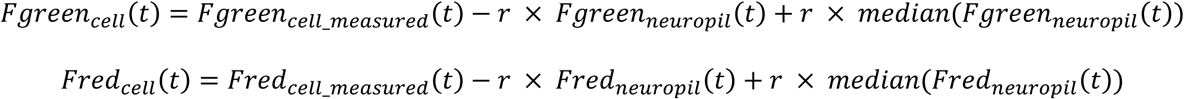

The ratio R(t) was then calculated as:

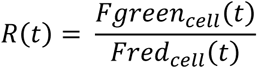

Slow timescale changes were removed by subtracting the 8^th^ percentile of a moving 14 s temporal window from R(t). ΔR/R_0_ was calculated as:

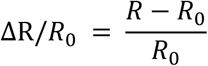

where R_0_ was measured over a 1 s period before the visual stimulation as the median of the individual mean baseline ratio signal of each trial. Visual responses were then extracted from trial-averaged responses as mean fluorescence ratio change over the full stimulus interval.

To determine visual responsiveness, a one-way ANOVA was performed over all averaged stimulation trials per orientation as well as R_0_ periods for each eye in the case of monocular stimulation. For binocular stimulation, a one-way ANOVA was performed over all averaged stimulation trials per condition as well as R_0_ periods. In both cases, neurons with *p*-values < 0.05 were identified as visually responsive.

Orientation-tuned cells were defined as neurons that showed a significant difference in responsiveness (p < 0.01, one-way ANOVA) over all presented grating directions in the ipsilateral, the contralateral or both eyes. The calculation of stimulus selectivity was performed on eye-specific responses that were significant in 50 % of the trials of at least one stimulus direction of a single eye exposure.

OD was determined by the OD index (ODI) for each individual cell:

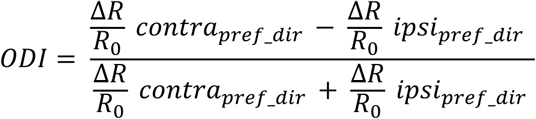

Where an ODI value of 1 or −1 displays exclusive contra- and ipsilateral dominance, respectively. Global orientation selectivity index (gOSI) was computed as 1 - circular Variance (circ. Var.):

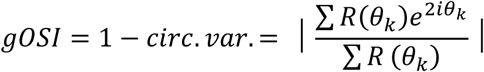

and global direction selectivity index (gDSI) was computed as:

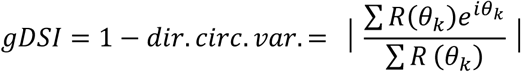

*R*(*θ*_*k*_) is here the mean response to the direction angle (*θ*_*k*_). Perfect orientation/direction selectivity is indicated with gOSI/gDSI of 1, whereas a gOSI/gDSI value of 0 indicates no orientation or direction selectivity. The preferred orientation and direction were computed by fitting a double-Gaussian tuning curve to the data. For binocular cells, the preferred orientation was defined as that one from the dominant eye, as determined by the sign of the ODI.

To determine spontaneous activity events in the dark, the baseline (R_0_) was calculated by taking the 8^th^ percentile of a 20 s moving window across the entire spontaneous activity period, and averaging these values. Then this R_0_ was used in the same way as the one described above for the visual stimulation protocols to yield ΔR/R_0_. Calcium event detection was performed by first taking the derivative of the low passed calcium trace (cut-off at 5 Hz). An event onset was defined as any point where the z-scored trace crossed a value of 2.

Population coupling was estimated by the correlation of a cell’s ΔR/R_0_ trace to the average ΔR/R_0_ trace of the rest of the population within the same recording. The population values were z-scored within each recording to compare data across multiple experiments. Finally, noise correlations were calculated by first computing the Pearson pairwise correlation matrix of aligned single trial responses between cells. Then, for each cell, the vector of its correlation values was taken with every other cell and averaged (excluding the target cell). Since the eyes were stimulated separately, for each cell only the average coming from its preferred eye stimulation sequence was considered. There were no differences when taking the average of the correlations versus when taking the noise correlations for the preferred direction stimulus (data not shown).

### Photostimulation analysis

The spatial resolution of LSPS by UV glutamate uncaging was calculated based on the size of the excitation profiles as the mean weighted distance from the soma (d_soma_) of AP generating sites using the following equation:

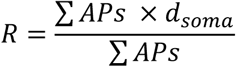

LSPS by UV glutamate uncaging induces two types of responses: 1) direct glutamate uncaging responses originating from direct activation of the glutamate receptors of the recorded neuron. 2) synaptically mediated responses originating from the activation of presynaptic neurons (Supplementary Fig. 2). Responses to the LSPS stimulation protocol (both for EPSCs and IPSCs) were quantified in the 150 ms window following the uncaging light-pulse, since this is the time window were evoked activity is normally observed. Considering the diversity of responses encountered in these experiments, a heuristic analysis scheme was devised to address the main observed cases:

1. Inactive traces were excluded by only considering those responses with a deflection higher than 2 S.D. over the baseline at any point. Additionally, traces that only had a significant response in one (out of x) repetition were also excluded.
2. Then, purely synaptic responses, i.e. those that correspond only to activation of the presynaptic terminal via uncaged glutamate - the ones of interest in this study - were selected by taking the traces that passed the 2 S.D. threshold only after a 7 ms window from the offset of stimulation.
3. For responses that did not pass the previous criterion, inspection by eye indicated that several of them presented all the identifiable features of purely synaptic responses but seemed to cross the threshold slightly earlier than 7 ms. An additional set of experiments performed on a subset of cells, where maps were measured before and after application of TTX (and hence before and after only direct responses were present) were added to measure the effect of considering these intermediated cases (∼5% of the total number of traces). These experiments showed that by using a secondary window of 3.5 ms, the error incurred in including these intermediate traces is ∼20 % (data not shown). Therefore, this secondary window was used to include a second batch of traces into the synaptic response pool.
4. Finally, those traces that did not pass the secondary window were then blanked, and a 4-dimensional interpolation method (via the MATLAB function “griddatan”) was used to infer their temporal profiles based on their 8 neighboring pixel activities in space and time. In the aforementioned TTX experiments (data not shown) every position with a response was observed to have a synaptic component, but the addition of this synaptic component and the overlapping direct component is non-linear. Therefore, this interpolation method was used to extract the synaptic component partially masked in the raw traces by the direct response. The approach relies on the observation that the synaptic responses of neighboring positions are similar across time, therefore indicating that information on the synaptic responses masked by direct responses is contained in the responses surrounding them. These interpolated responses were then incorporated into the maps as synaptic responses, and used in all subsequent calculations and figures. For excitatory input maps, the first two stimulation rows were excluded since excitatory input from L1 originated from cells in L2/3-L5 having apical tuft dendrites in L1, which fired action potentials when their tufts were stimulated in L1 (see Supplementary Fig. 3).

### Centroid calculation

To calculate the weighted centroid for each map, the layers of interest (2/3, 4 or 5) were separated, the weighted centroid was calculated according to the following formula and then the centroid coordinates were translated from image coordinates to input map coordinates.

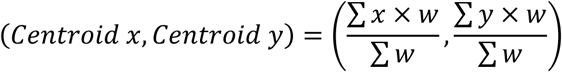

Here *x* and *y* are the horizontal and vertical coordinates of every pixel, and *w* is the value associated with that position. Then, all coordinates were translated to their anatomical location for further use and comparison with other cells. The distance between the cell soma and the centroid was utilized in most of the analyses.

### Input Map Principal Component Analysis (PCA)

Since the goal was to assess the main sources of variance driving the shape of the input distributions, before PCA the input maps were aligned based on the soma position of each cell. This involved shifting the maps vertically an integer number of stimulus rows until all the somata were in the same row. Then, all maps were internally normalized and used as features for PCA. The first 3 Principal Components for each input map were extracted (carrying roughly 60% of the variance in the dataset). PCA was performed separately for excitation and inhibition maps to keep their independence unbound. As a control, the combined excitation-inhibition PCA decomposition was calculated. For this, the feature vectors from excitation and inhibition for each map were concatenated, yielding a 512 element feature vector that was then used for the decomposition in the same way as described above.

### UMAP embedding

Uniform Manifold Approximation and Projection was utilized to visualize the distribution of different properties across the data on a cell by cell basis. The computational details of UMAP are described elsewhere (McInnes, Healy, and Melville 2018). Briefly, UMAP embeds data points from a high dimensional space into a 2D space preserving their high dimensional distances in a neighborhood. This permits effective visualization of the connections between data points. A UMAP implementation in Matlab developed by Meehan, Meehan and Moore (https://www.mathworks.com/matlabcentral/fileexchange/71902) was utilized. The excitatory input fraction from L4, the x and y coordinates of the inhibitory L2/3 input map centroid and the pial depth of the cell were used as the embedding parameters. 15 was used as the number of neighbors and 0.1 as the minimum distance (default parameters). The embedded points were color-coded depending on the property, either in a normalized scale or in a periodic scale when visualizing angles.

### Morphological reconstruction and analysis

The reconstruction of dendritic cell morphology was performed manually using the Simple Neurite Tracer of ImageJ. Reconstructions were quantitatively analyzed in MATLAB and with the open-source TREES toolbox (Cuntz et al. 2011). For Sholl analysis, the number of intersections between dendrites and concentric spheres centered on the soma was determined at increasing distances from soma (20 μm increments).

### Statistics

Data are reported as mean ± standard error of the mean (SEM). Correlation coefficients were calculated as Pearson’s correlation coefficient. The Circular Statistics Toolbox developed by Philipp Berens was utilized for circular correlation calculation (https://www.mathworks.com/matlabcentral/fileexchange/10676-circular-statistics-toolbox-directional-statistics).

Before comparison of data, individual data sets were checked for normality using the Kolmogorov-Smirnov Goodness-of-Fit test. None of the data sets considered in this study was found to be normally distributed. Therefore, paired or unpaired nonparametric statistics (Wilcoxon rank sum test, signed-rank or Kruskal-Wallis test on ranks with Bonferroni’s post hoc test for multiple comparison) were used for comparison. Two-tailed tests were used unless otherwise stated. Asterisks indicate significance values as follows: *p<0.05, ** p<0.01, *** p<0.001.

## Acknowledgements

We are grateful to Volker Staiger for cell tracing as well as technical support and to Michael Myoga for helping to build the *in vitro* setup. This study was supported by the Max Planck Society and the Deutsche Forschungsgemeinschaft (CRC 870; V.S., T.B., T.R., and M.H.).

## Author Contribution

S.W. and V.S. conceived the project, with input from M.H. and T.R.. S.W. planned and performed all experiments. D.G.N., T.R. and S.W. wrote the advanced analysis tools and D.G.N., S.W. and V.S. analyzed the data. S.W. and V.S. implemented LSPS at the patch-clamp setups. T.R. designed and built the *in vivo* 2-photon setup and developed the viral construct. S.W., D.G.N, and V.S. wrote the paper. T.R., M.H. and T.B. edited the paper. T.B. provided research environment.

