## Supplementary Figures for "Relationship between input connectivity, morphology and orientation tuning of layer 2/3 pyramidal cells in mouse visual cortex"

**Supplementary Material**

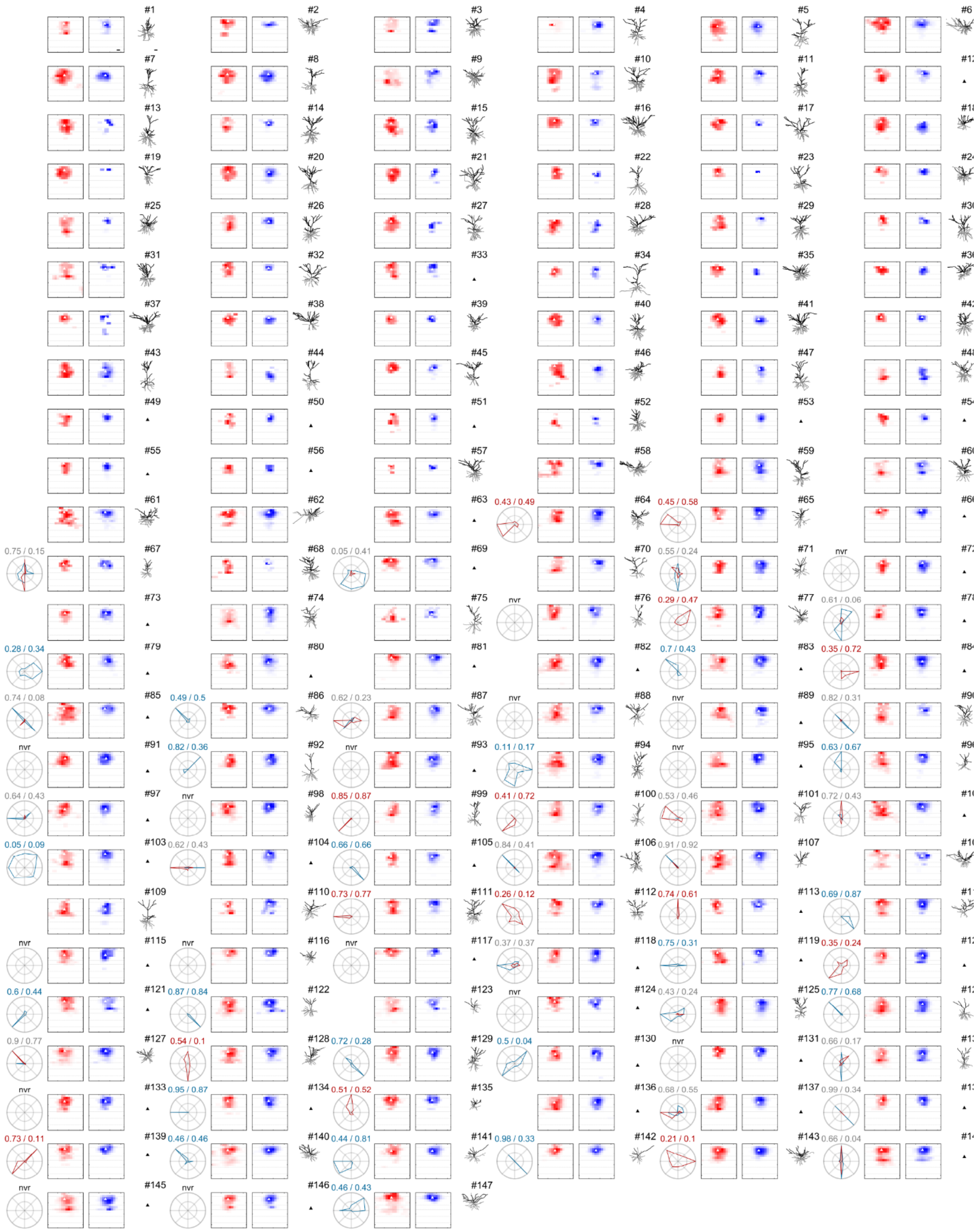

**Supplementary Figure 1:** Overview of complete data set of visual response properties, excitatory/inhibitory laminar input patterns and dendritic morphology for 147 L2/3 pyramidal cells.

The experimental ID is indicated at the top right for each cell. Data for each cell is displayed as follows: Left, polar plot displaying the eye-specific orientation and direction tuning of each cell, if data is available (0 degrees to left, angles increasing clockwise, as in polar plot in Fig. 1C). For binocular cells, both contra- (blue) and ipsilateral (red) eye responses are displayed, for monocular cells only that of the responsive eye. Eye-specific global orientation and direction selectivity indexes (gOSI, gDSI) are indicated at top. Middle, normalized excitatory and inhibitory input maps (scale bar: 100 µm). Right, dendritic morphology, if available. Apical tree in black, basal dendrites in grey (scale bar: 50 µm).

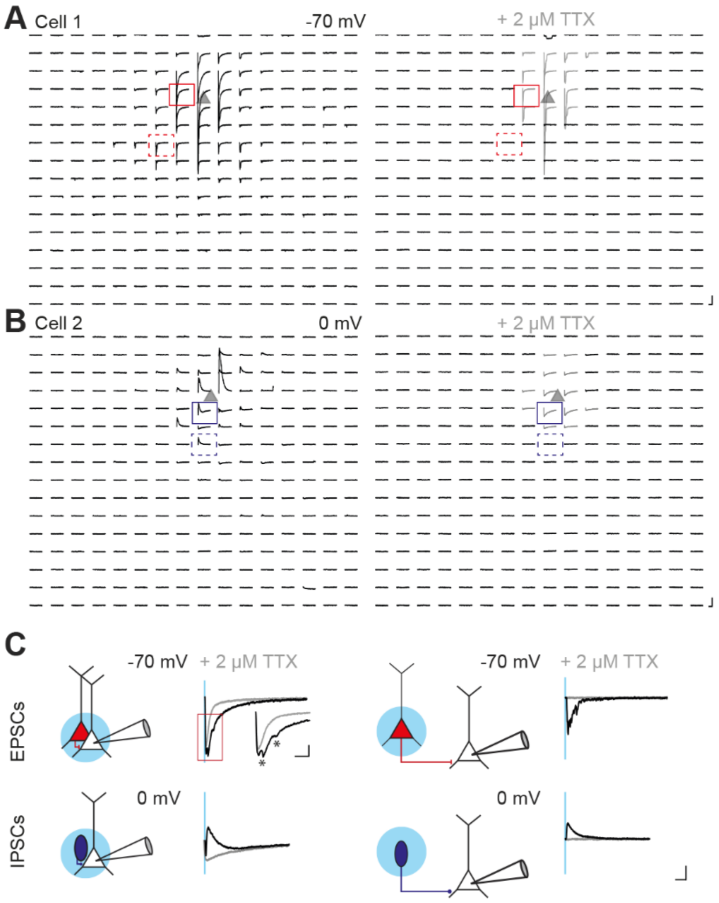


**Supplementary Figure 2:** LSPS response evaluation

**A** Traces from LSPS evoked EPSCs (at -70 mV) measured at 16 x 16 locations from a L2/3 PC before (left) and after infusion of 2 µM TTX (right, scale bars: 250 pA, 100 ms). Boxes highlight example traces proximal (solid box) and more distal (dashed box) to the soma, see C. **B** Same as A for a cell clamped at 0 mV to measure LSPS evoked IPSCs. **C** Schematic illustrating different stimulation scenarios giving rise to pure (right) and synaptic responses contaminated by direct stimulation (left). Large glutamate responses evoked directly in the recorded neuron (direct response) obscure the evoked responses arising from connected cells in the immediate vicinity (synaptic responses) when the laser spot covers the recorded cell (left, black trace, taken from cell 1 in A, left panel, highlighted with solid box). Direct glutamate responses are isolated by blocking photostimulation-evoked synaptic currents with TTX (grey trace, taken from cell 1 in A, right panel, highlighted with solid box). Inset, traces within box on extended timescale. Asterisks mark synaptic responses. When a cell is clamped at 0 mV, the direct response is smaller compared to the synaptic response (bottom, taken from cell 2 in B, highlighted with solid box). Synaptic and direct responses can be distinguished by their temporal delay. Because the onset of the direct responses is locked to the start of the laser pulse (blue lines) they can be detected using a temporal window between photostimulus and onset of the synaptic responses (7 ms, see Methods). Examples of photostimulus-evoked pure excitatory (top right) as well as inhibitory (bottom right) synaptic responses arising from more distal presynaptic neurons (from dashed boxes in A and B). The synaptic responses occur with a temporal delay after the photostimulus. The evoked synaptic currents (black) are blocked after TTX infusion (grey) without revealing a direct component (scale bars: 250 pA, 100 ms).

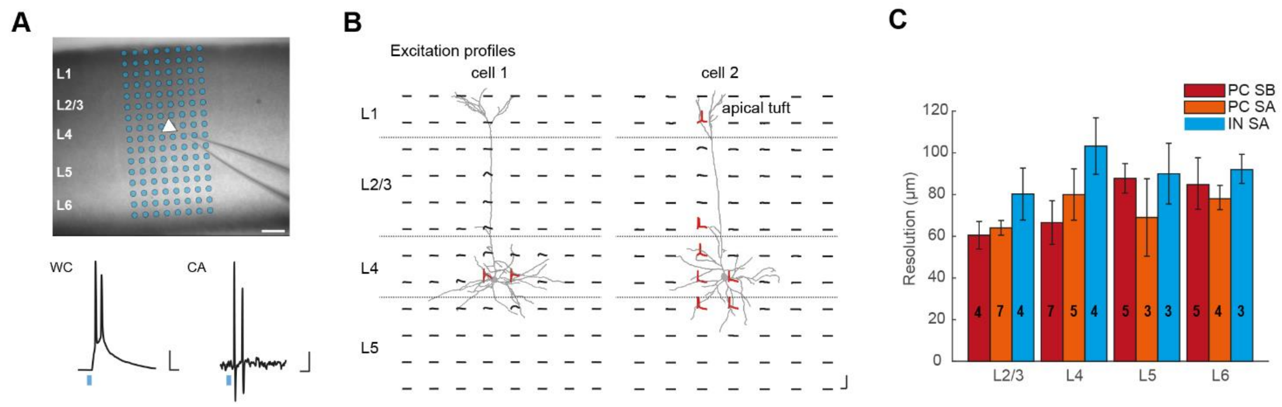


**Supplementary Figure 3:** Spatial resolution of photostimulation across cortical layers and cell types.
**A** Example L4 pyramidal cell recorded for assessing spatial resolution. The image of the bV1 slice is superimposed with the stimulation grid (50 µm spacing, scale bar: 100 µm, top). Photostimulation evoked response either in whole-cell (WC, left) or cell-attached (CA, right) configuration (bottom). Blue box indicates photostimulus (scale bars: right, 2 mV, 20 ms; left, 0.2 mV, 20 ms). **B** Voltage traces for each stimulus spot of two L4 pyramidal cells recorded in WC current clamp. Generally, APs occur only close to the soma of the recorded cell (left). Neurons with apical tufts in L1 can also fire an AP upon stimulation there (right). For this reason and because of the absence of excitatory cells in L1, we excluded stimulation sites in layer 1 synaptic input maps (see Methods, scale bars: 100 pA, 100 ms). **C** Spatial resolution of LSPS evoked action potential generation in PCs and interneurons (INs) in the different cortical layers for setup A (SA) and setup B (SB, see Methods). Spatial resolution of photostimulation is measured as the mean weighted distance from the soma of AP generating stimulation sites. The numbers of recorded neurons are displayed within the bars.

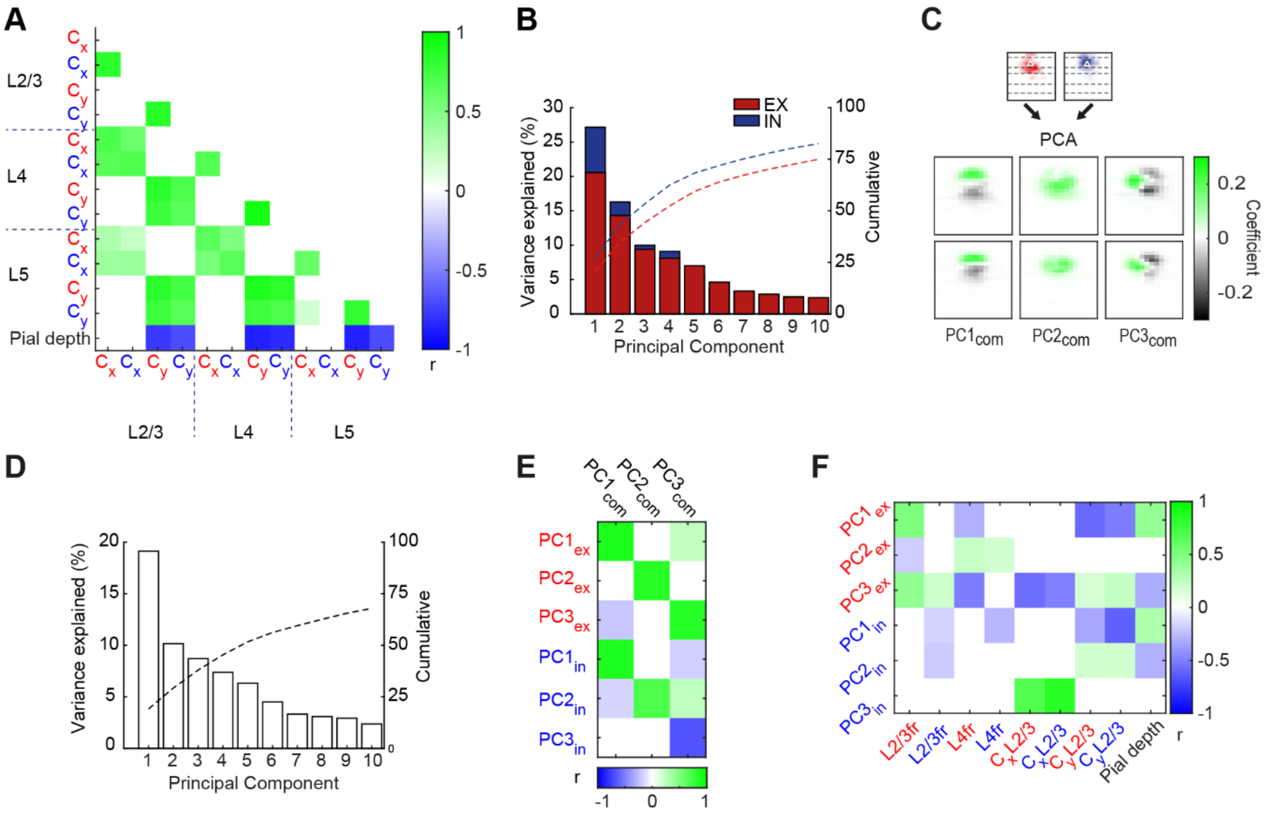


**Supplementary Figure 4:** Principal component analysis reveals relevant features in excitatory and inhibitory maps. **A** Correlations between pial depth, $C_x$ and $C_y$ within L2/3, L4 and L5 for excitation and inhibition. Colors indicate the Pearson correlation coefficient between the pair of parameters according to the color bar on the right. Coefficients with p values > 0.05 are set to 0 ($C_y$ vs. pial depth, excitation, L2/3: r=-0.84, p<0.001; L4: r=-0.87, p<0.001; L5, r=-0.83, p<0.001; inhibition, L2/3: r=-0.81, p<0.001; L4: r=-0.84, p<0.001; L5: r=-0.74, p<0.001, n=147, Pearson's correlation coefficient). **B** Percentage of variance explained by each of the first 10 principal components for excitation and inhibition. Lines show the cumulative sum of the variance explained. **C** Principal component analysis (PCA) using the combined 16x16 normalized excitatory and inhibitory input maps (top). Before PCA, the input maps were vertically and horizontally aligned (see Methods). Input maps of the first three principal component eigenvalues (bottom, compare to Fig. 3D). **D** Percentage of variance explained by the first 10 principal components for combined excitation/inhibition. Line shows the cumulative sum of the variance explained. **E** Correlations between the first three principal components for combined and separate excitation and inhibition. **F** Correlations between the first three principal components for excitation and inhibition and extracted input map features displayed in Figures 2 and 3. Colors indicate the Pearson correlation coefficient between each pair of parameters according to the color bar on the right. Coefficients with p values > 0.05 are set to 0. Colors indicate the Pearson correlation coefficient between the pair of parameters according to the color bar at the bottom. Coefficients with p values > 0.05 are set to 0.

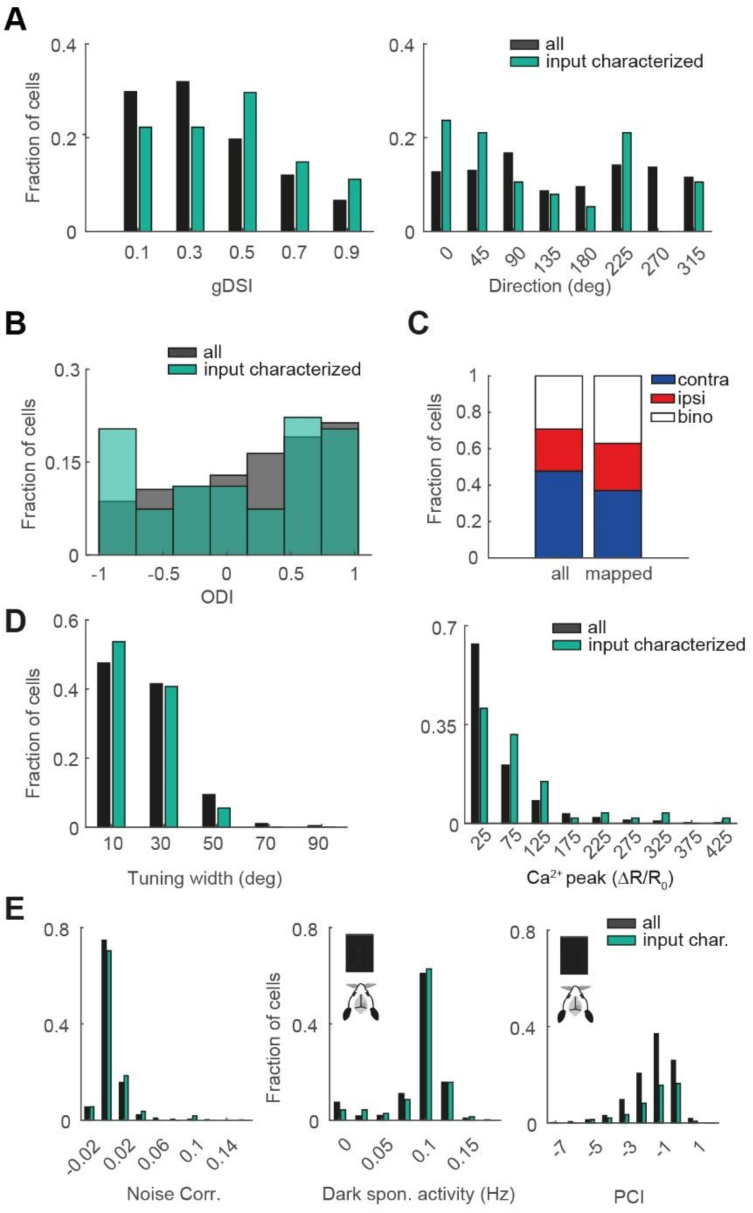


**Supplementary Figure 5:** Response properties of L2/3 pyramidal cells in bV1.

**A** Global direction selectivity index (gDSI) and direction preference (DIR) distributions of all *in vivo* characterized cells (black, n=1134, n=693) and subset of cells that were *in vivo* / *in vitro* characterized (green n=54, n=38). For the DIR distributions, only cells with gDSI>0.25 are displayed. **B** Same as A for ocular dominance (ODI, n=1134, n=54). **C** Fraction of contra-, ipsilateral and binocular responsive cells for all *in vivo* characterized cells (n=541, n=262, n=331) and subset of cells that were *in vivo* / *in vitro* characterized (n=20, n=14, n=20). **D** Same as A for tuning width (left) and $Ca^{2+}$ peak amplitude (right). **E** Same as in A for noise correlations (left), spontaneous activity events (middle) and population coupling index (PCI, right n=2237, n=70). Note that spontaneous activity and PCI were measured in complete darkness.

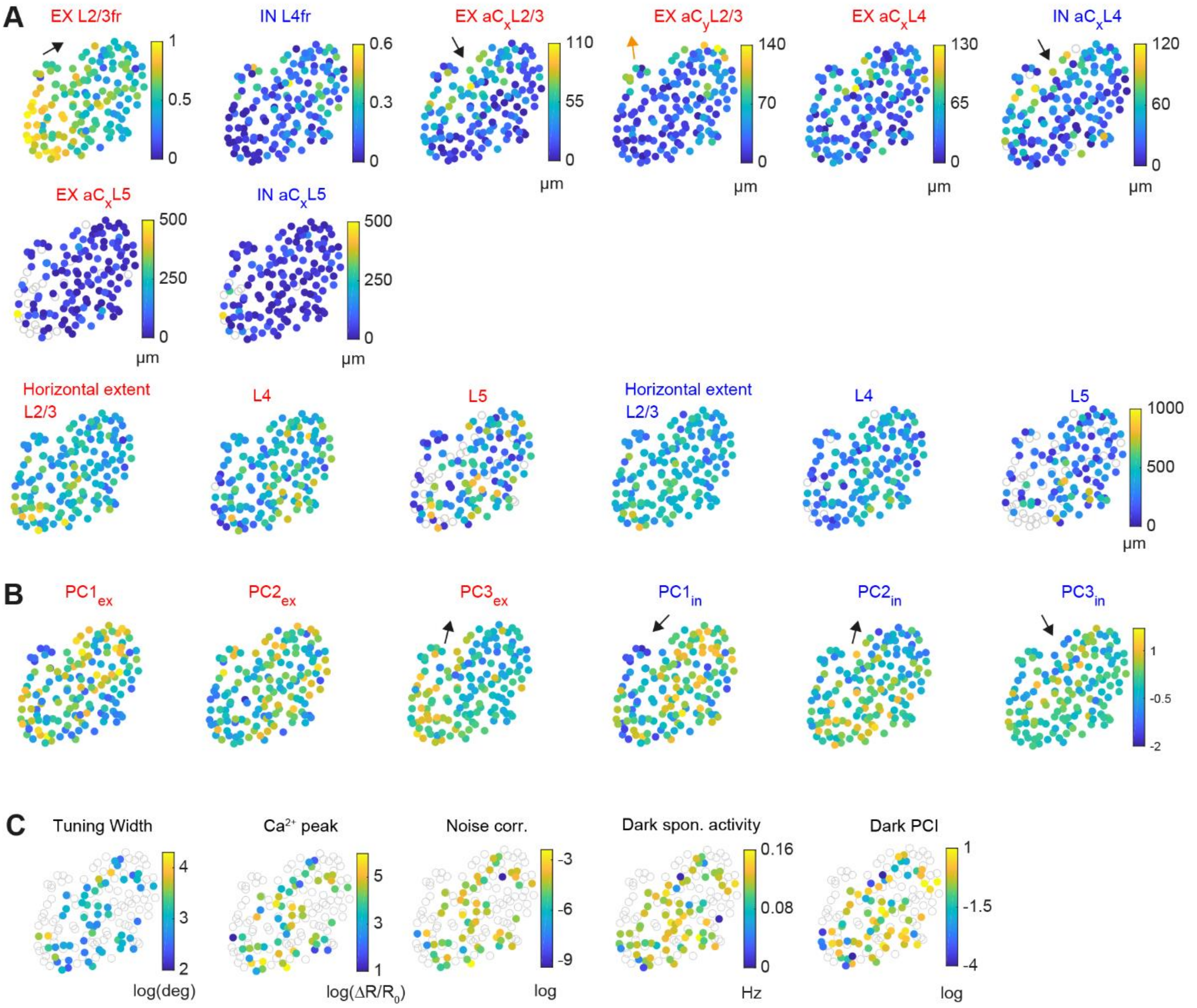


**Supplementary Figure 6:** UMAP projections of input map features, principal components and *in vivo* response properties.
**A** UMAP projections of different input map features. The same embedding as in Fig. 4 was used. **B** UMAP projections of the first three excitatory and inhibitory principal component weights. **C** UMAP projections of tuning width, $Ca^{2+}$ peak amplitude, noise correlations, spontaneous activity events and population coupling index. Note that some parameters are displayed with a log scale for better visualization.

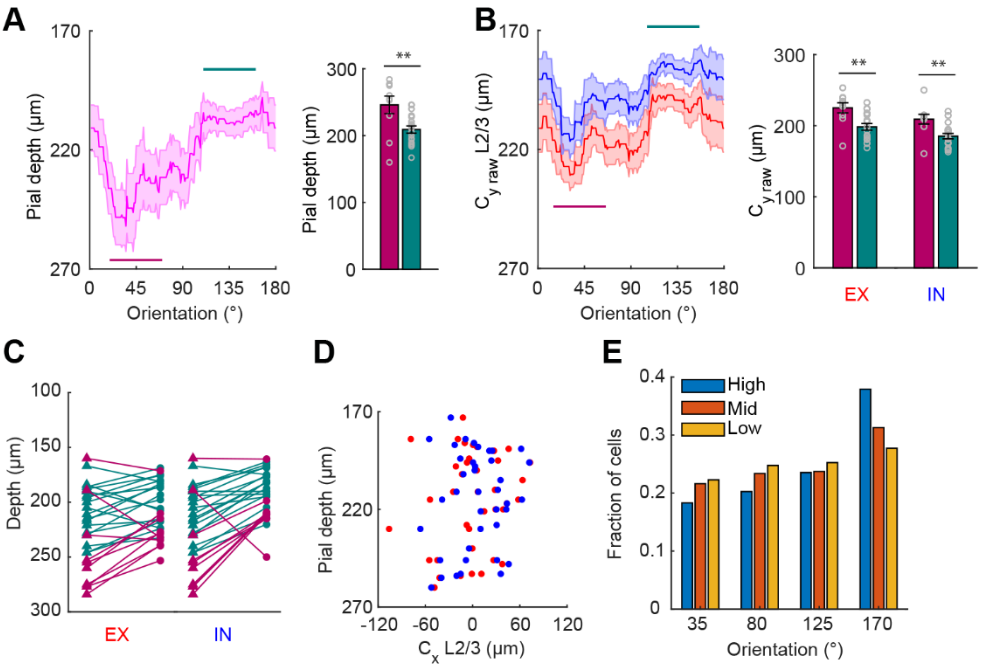


**Supplementary Figure 7:** Relationship between the relative centroid positions ($C_x$, $C_y$) and other cell properties.
**A** Rolling average of the pial depth as a function of preferred orientation (left, mean ± SEM, n=50), and pial depth comparison between the two angle ranges indicated (right, n=10, n=18). One-tailed Wilcoxon rank-sum**. B** Same as in A for the the raw vertical centroid position $C_{y\,raw}$ , for excitation (red) and inhibition (blue). Averages of $C_{y\,raw}$ for the indicated angle ranges (right). **C** Total depth of a cell's soma (triangle) compared to the depth of its centroid (circle) plotted for both excitation and inhibition. Turquoise and purple indicate the 125 and 35° angle ranges, respectively. **D** Pial depth of the soma plotted against the horizontal distance to centroid for both excitation (red) and inhibition (blue). **E** Distribution of preferred orientations of all cells characterized *in vivo* with increasing depth in L2/3 (high: 100-200 µm, n=153; mid: 200-300 µm, n=569; low: 300-400 µm, n=191, N=31).

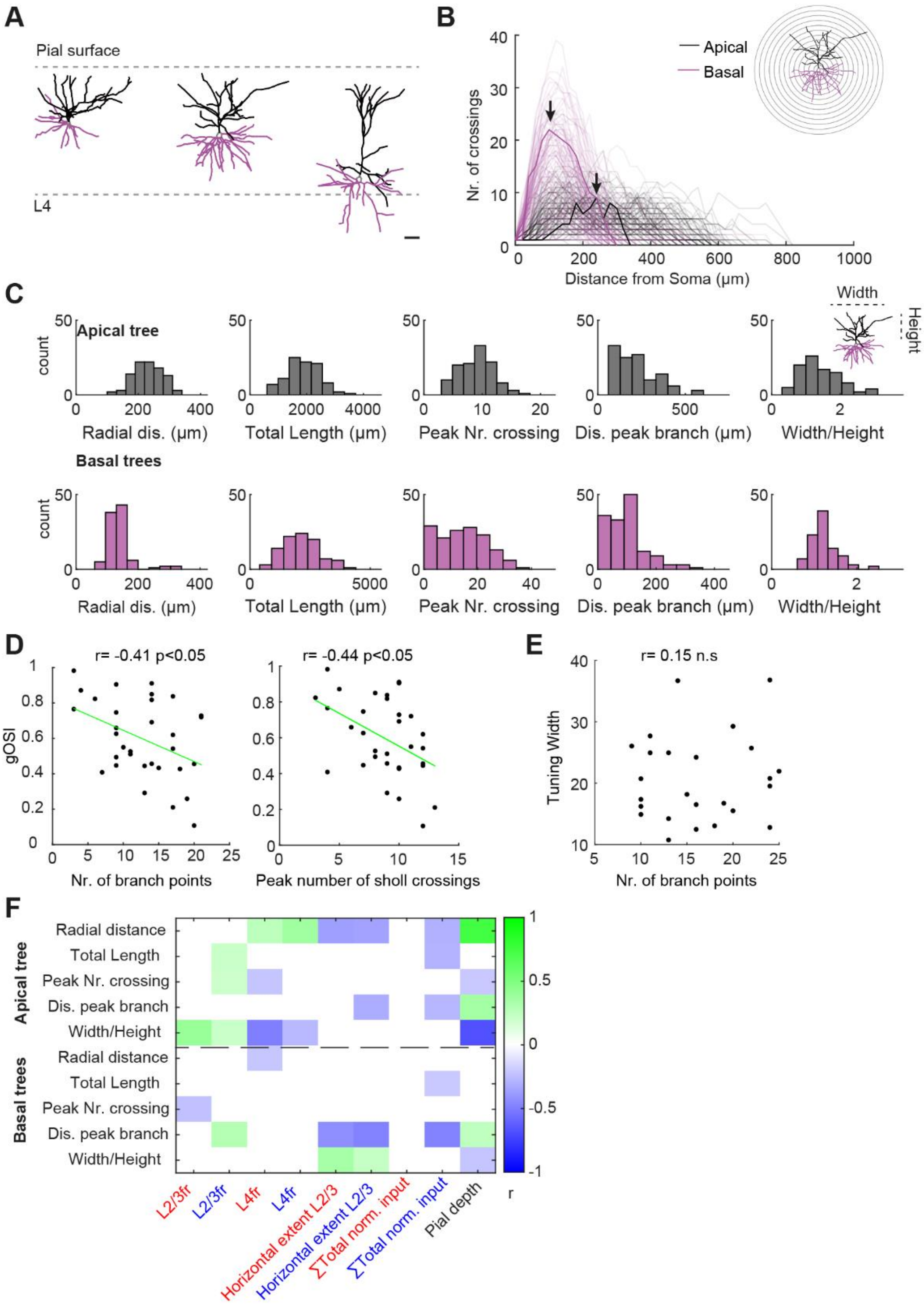


**Supplementary Figure 8:** Morphological features of L2/3 pyramidal cells and their relation to input map properties. **A** Reconstructed dendritic morphology of cells in the upper, middle and lower part of L2/3 (scale bar: 50 µm). Apical dendrites, black, basal dendrites, purple. **B** Sholl analysis for apical and basal dendrites (n=97). The number of crossings was determined using concentric spheres centered around the soma with 20 µm increments. Bold lines refer to the example cell in inset. Arrows indicate the peak number of crossings. **C** Distributions for five morphological features for apical (top) and basal dendrites (bottom). **D** gOSI plotted against the number of branch points (left) and the peak number of Sholl crossings (right) of the apical tree. **E** Tuning width plotted against the number of branch points of the basal trees. **F** Correlations between morphological features for apical and basal trees and input map features (n=97). Colors indicate the Pearson correlation coefficient between the pair of parameters according to the color bar on the right. Total length, maximal extent and distance of peak branch (Dis. peak branch) are in µm. Coefficients with p values > 0.05 are set to 0.
